# From Attraction to Repulsion: Omicron-Driven Rewiring of Nanobody Interfaces on the SARS-CoV-2 RBD

**DOI:** 10.1101/2025.07.18.665531

**Authors:** Mert Golcuk, Fareeda E. Abu-Juam, Derman Basturk, Ayten Dilara Gursel, Clara Xazal Buran, Reyhan Metin Akkaya, Ahmet Yildiz, Mert Gur

**Author notes:** Corresponding Author: Mert Gur.

## Abstract

Mutations in the SARS-CoV-2 spike receptor-binding domain (RBD) of the Omicron variant enable broad escape from neutralizing antibodies (Abs) and nanobodies (Nbs), yet the atomistic basis of epitope-dependent loss of Nb binding remains unclear. We performed all-atom molecular dynamics (MD) simulations of 13 Nbs bound to the Omicron RBD to characterize their binding modes, binding-pose stability, and interfacial dynamics. Analysis of the trajectories and corresponding free-energy landscapes revealed that most Nbs occupied a single dominant basin but showed varying degrees of within-basin heterogeneity and often exhibited orientation shifts, whereas two Nbs (Nb21 and Sb14) showed pronounced pose plasticity and dissociated from the RBD. For the remaining 11 Nbs, covariance-matrix analysis mapped stabilizing and destabilizing residue couplings, revealing recurrent hydrophobic anchor patches within the receptor-binding motif and Nb-specific interaction patterns reflecting differences in the sequences and chemistries of their complementarity-determining regions. Omicron substitutions rewired interaction networks, shifted Nb binding orientations relative to the corresponding experimentally determined reference structures (obtained with the ancestral WT RBD for all Nbs except H3), and introduced unfavorable repulsion, either directly at mutation sites or indirectly via mutation-driven reorientation, thereby reducing interfacial stability. Low-speed steered MD unbinding simulations of the 11 Nb–RBD complexes and the ACE2–RBD complex, performed at a pulling velocity within the range used in high-speed AFM-based force spectroscopy, yielded lower mean unbinding work for all 11 Nbs than for ACE2. These simulations also revealed heterogeneous, stepwise Nb unbinding pathways in which receptor-binding ridge (RBR)-mediated anchoring maintained partial Nb attachment through newly formed interactions and stabilized transient intermediate binding poses. Together, these Nb–RBD interaction fingerprints and pathway-resolved unbinding simulations pinpoint epitope-specific determinants and mutation-induced clash sites, providing a mechanistic basis for diminished Nb binding.

## INTRODUCTION

Nanobodies (Nbs) are single-domain antibody (Ab) fragments that are approximately an order of magnitude smaller (∼12–15 kDa) than full-length Abs while offering high stability and comparable affinity.^1,2^ Their compact, monomeric structure and often elongated complementarity-determining region 3 (CDR3) loop enable Nbs to bind cryptic epitopes on target proteins that are frequently inaccessible to conventional Abs.^3^ Since the emergence of SARS-CoV-2, various Nbs that target the spike glycoprotein’s receptor-binding domain (RBD) have been identified to neutralize infection by blocking receptor engagement or by destabilizing the RBD–ACE2 interface (**Figure 1A**).^4,5^ For example, Nb H11-H4 can bind the RBD and effectively displace ACE2 via electrostatic repulsion without direct overlap, whereas Nb Ty1 binds the RBD at the ACE2 site, competing directly with the receptor.^4^ Such mechanistic differences illustrate the diversity of mechanisms by which Nbs neutralize the virus and underscore the importance of mapping RBD– Nb interactions as dynamic, residue-resolved interfaces rather than as static epitope overlaps alone.

**Figure 1.**
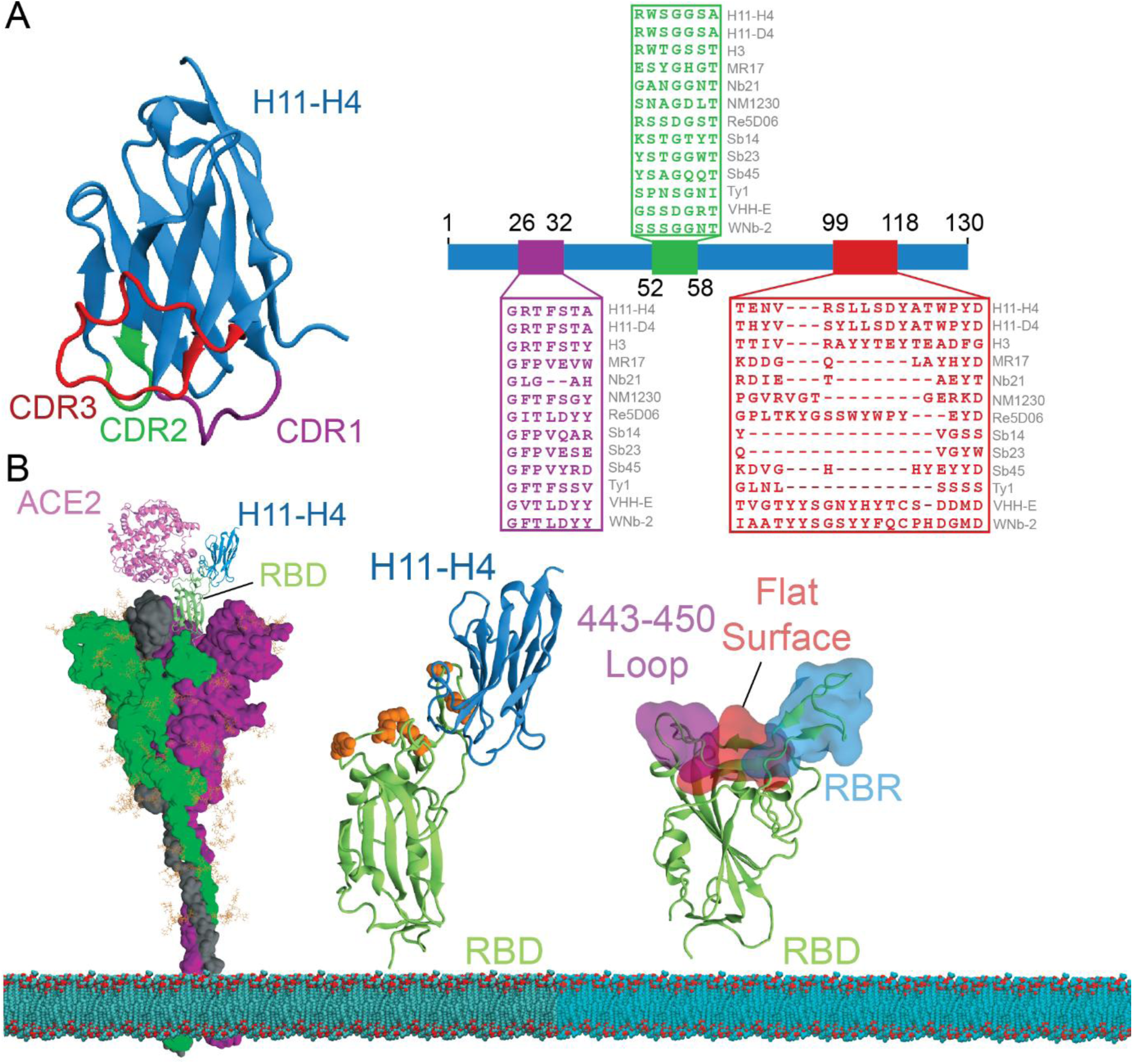
Nanobody CDR architecture and overlap with the Omicron RBD–ACE2 interface. *(A)* Cartoon representation of the Nb (blue) variable domain with CDRs highlighted: CDR1 (residues G26–A32 for Nb H11-H4, magenta), CDR2 (residues R52–A58 for Nb H11-H4, green), and CDR3 (residues T99–G118 for Nb H11-H4, red). Nb H11-H4 is shown as a reference example, and the right panel shows Clustal Omega^48^ multiple sequence alignments (aligned by the entire Nb sequence) of CDR1–3 from H11-H4 and other Nbs, illustrating the typically greater sequence diversity and length variability of CDR3^49^ compared with CDR1 and CDR2. (B) Structural model of the Omicron SARS-CoV-2 spike trimer (surface) with the RBD (green) bound by Nb (blue) and superposed with ACE2 (pink), showing their relative positions on the RBD. The lipid bilayer is shown at the bottom. Center, close-up of the H11-H4–RBD complex (RBD in green) with highlighted Omicron mutation sites (T478K, E484A, Q493R, Q498R, N501Y) shown as orange spheres. Right, surface representation of the RBD highlighting the RBR (blue), the S443–N450 loop (purple), and an adjacent flat surface (salmon).

SARS-CoV-2 continues to evolve, accumulating mutations that enable emerging variants to evade existing Abs and sustain viral circulation. The Omicron lineage, in particular, carries a large number of mutations, many clustered in the spike RBD (**Figure 1B**).^6–8^ Because this domain mediates spike engagement with ACE2, a key step in spike-dependent viral entry, it has been a principal antigenic target for vaccine design, monoclonal Abs (mAbs), and Nbs.^9–13^ Within the RBD, the receptor-binding motif (RBM; approximately S438–Y508, depending on the boundary definition) forms most of the ACE2-contacting surface. The spike RBM is commonly described as comprising three structural elements: (i) the receptor-binding ridge (RBR; ∼T470–C488), (ii) the S443–N450 loop, and (iii) the relatively flat intervening surface **(Figure 1B)**.^14^ In an alternative, largely overlapping nomenclature, the ACE2-contacting interface is partitioned into contact regions 1–3.^15^ Omicron substitutions are distributed across all of these RBM sub-elements, placing immune escape mutations directly within or adjacent to many Ab/Nb epitopes.

Consistent with ongoing antigenic drift, many mAbs and Nbs that were highly effective against the wild-type (WT) strain have progressively lost potency with the emergence of the Delta and Omicron variants.^8,16–18^ In fact, the neutralizing effect of >85% of tested RBD-targeted Abs was diminished by Omicron mutations.^19,20^ This has reduced the clinical efficacy of several Ab therapies and led to the revocation of some emergency-use therapeutic authorizations (EUAs).^21^ Therefore, each emerging variant may necessitate new or re-engineered Abs and Nbs, and rapid, systematic evaluation of Ab/Nb–antigen interactions. Conventional experimental screening alone is not able to keep pace with the rate of SARS-CoV-2 antigenic drift. All-atom molecular dynamics (MD) simulations have become a valuable tool for rapidly assessing how RBD mutations impact Ab/Nb binding.^4,16–18^ By sampling protein–protein interfaces on ns to μs timescales, MD simulations can reveal how mutations perturb local environments, interfacial contacts, interaction networks, and binding-pose stability. Across computational studies of SARS-CoV-2 Ab and Nb recognition, MD has been used to resolve residue-level interaction hotspots, trace mutation-driven rewiring of interfacial contacts, and explain why some binders retain activity whereas others lose binding as new variants emerge.^17,18^ Within this broader context, our previous Nb studies used MD simulations to elucidate the binding mechanisms and relative binding strengths of Nbs against earlier variants and to explore how Nb binding can destabilize and displace bound ACE2 from the RBD surface,^4,16^ thereby providing insight into how mutations in Alpha, Beta, and Delta variants alter Nb efficacy.

At the interface level, the RBD–Nb bound state often comprises an ensemble of closely related substates rather than a single fixed pose, with one substate corresponding to the most populated binding mode. Experimentally resolved or ML/AI-predicted structures usually represent one member or reference model of this ensemble and therefore may not capture less populated motions, transient contacts, or mutation-induced reorientation events. MD simulations can sample the conformational space accessible to bound complexes over the simulated timescale, and principal component analysis (PCA) provides a powerful way to reduce these trajectories to the dominant collective motions (principal components, PCs) of the binder relative to the interface.^22–27^ By projecting sampled conformations onto the top PCs and generating free energy surfaces, dominant bound basins, alternative substates, and their relative populations can be identified within the bound ensemble.^25–29^

Yet, resolving the full spectrum of stabilizing and destabilizing residue-residue interactions within individual bound substates or across the complete bound ensemble remains challenging. Repulsive contacts, including same-charge side-chain encounters and hydrophobic–charged polarity mismatches, can be short-lived because they drive local separation after a transient approach. Hydrophobic contacts pose a complementary challenge: a simple distance cutoff can over-assign stabilizing nonpolar contacts when packing is loose, water remains between the two surfaces, or nearby charged and polar groups disrupt the interaction. Similarly, solvent exposure, ionic screening, or side-chain orientation can weaken an apparent repulsive interaction. To address this limitation, we recently developed a covariance matrix–based method^30^ that efficiently extracts stabilizing (attractive) and destabilizing (repulsive) interactions governing RBD–Nb binding. Rather than classifying all close contacts solely by geometric criteria, such as distance and angle, this workflow first uses correlations from MD trajectories to prioritize dynamically coupled residue pairs. Unlike static structures or purely geometric contact maps, these correlations are computed from motions generated in the fully simulated environment, and therefore implicitly reflect the net dynamical effects of attractive and repulsive contacts, solvation, ionic screening, steric packing, and side-chain rearrangements. The prioritized correlated pairs are then assigned interaction types using chemical identity and geometric criteria.

Bound-state analyses provide critical information, but they do not fully describe how an interface fails during detachment. Mechanical resistance to dissociation is governed not only by the endpoint free-energy difference between the bound and unbound ensembles, but also by the free-energy barriers encountered along the unbinding pathway. These barriers can exceed the endpoint difference and are shaped by intermediate binding poses, pathway heterogeneity, sequential contact-rupture events, and local reorientations during unbinding. Steered molecular dynamics (SMD) simulations, the computational analogue of single-molecule force spectroscopy,^31^ have emerged as a widely used method to simulate the unbinding of proteins under force and quantify resistance to unbinding while sampling ensembles of unbinding pathways.^32,33^ At finite pulling speeds, SMD work should be interpreted as a relative measure of mechanical resistance to forced detachment, rather than as a direct equilibrium binding free energy. Nevertheless, the nonequilibrium-work values provide a physical basis for comparing detachment under identical pulling conditions, and, by the second law of thermodynamics, the mean work provides an upper bound to the corresponding unbinding free-energy difference; ⟨W⟩ ≥ ΔG_unbinding_.^32,34–37^ This makes SMD simulations particularly useful for Ab/Nb–RBD complexes, in which similar bound poses can detach through different barriers and interaction-rupture sequences that control dissociation.^4,16^

Here, we performed an extensive set of physics-based all-atom MD simulations (24.8 μs total simulation time), including both conventional MD and SMD, to investigate previously uncharacterized binding modes and interfacial dynamics of a panel of Nbs (H11-D4,^5^ H11-H4,^5^ H3,^38^ MR17,^39^ Nb21,^40^ NM1230,^41^ Re5D06,^42^ Sb14,^43^ Sb23,^44^ Sb45,^43^ Ty1,^45^ VHH-E,^46^ and WNb-2^47^) bound to the Omicron BA.2 spike RBD. We used PCA-derived free-energy surfaces from the conventional MD simulations to define the dominant bound substates^25–28^ for each RBD–Nb complex. For complexes that retained a persistent bound state, we then applied our covariance-based interaction-mapping method^30^ to map stabilizing and destabilizing interfacial interactions and quantify their persistence. We then used low-speed SMD unbinding simulations, with a pulling speed in the range of high-speed force spectroscopy experiments,^33^ to compare the resistance of selected RBD–Nb complexes and the RBD–ACE2 complex to forced detachment and characterize their unbinding pathways. This integrated workflow connects bound-state ensemble heterogeneity, residue-level attraction-repulsion fingerprints, and mechanical resistance to detachment, revealing that Omicron substitutions can shift Nb binding orientations, rewire interfacial interaction networks, and, where WT data are available,^4,16^ reduce resistance to unbinding relative to the corresponding WT RBD–bound complexes; all tested Omicron-bound Nbs also exhibited lower resistance to unbinding than ACE2. These dynamic, residue-resolved fingerprints identify variant-sensitive features of Nb recognition and provide a basis for engineering more resilient Nbs against emerging SARS-CoV-2 variants.

## METHODOLOGY

### MD Simulations of Nanobody–Omicron RBD Complexes

All-atom MD simulations were performed for each RBD–Nb complex to characterize binding dynamics. Simulations were performed on the SARS-CoV-2 Omicron BA.2 spike glycoprotein’s RBD, which includes the array of mutations characteristic of this variant. Initial atomic coordinates for the Omicron RBD–Nb complexes were obtained by introducing relevant point mutations into the known SARS-CoV-2 RBD–Nb complexes (**Table 1**). N-linked glycosylation at N343 on the RBD was included based on our previous studies.^8,16^ Each complex was placed in a cubic simulation box with 25 Å of padding in all directions using the explicit TIP3P water model, ensuring ∼50 Å separation between periodic images. The solvated system was neutralized, and the ion concentration was set to 0.15 M NaCl. The final system sizes ranged from 135,000 to 170,000 atoms (**Table 1**).

**Table 1.** SARS-CoV-2 RBD-Nb complexes and system sizes.

| Nanobody | PDB ID | System Size (atoms) |
| --- | --- | --- |
| H11-D4 | 6YZ5<sup>5</sup> | ~150,000 |
| H11-H4 | 6ZBP<sup>5</sup> | ~150,000 |
| H3 | 7OAQ<sup>38</sup> | ~135,000 |
| MR17 | 7C8W<sup>39</sup> | ~160,000 |
| Nb21 | 7MDW<sup>40</sup> | ~150,000 |
| NM1230 | 7B27<sup>41</sup> | ~140,000 |
| Re5D06 | 7OLZ<sup>42</sup> | ~170,000 |
| Sb14 | 7MFU<sup>43</sup> | ~140,000 |
| **Sb23** | 7A29<sup>44</sup> | ~135,000 |
| **Sb45** | 7N0G<sup>43</sup> | ~140,000 |
| **Ty1** | 6ZXN<sup>45</sup> | ~145,000 |
| **VHH-E** | 7KN5<sup>46</sup> | ~160,000 |
| **WNb-2** | 7LX5<sup>47</sup> | ~155,000 |

The Omicron RBD–Nb complexes were energy minimized for 10,000 steps to relieve steric clashes, followed by a multi-stage equilibration protocol. Initially, protein heavy atoms were restrained, and water and ion molecules were relaxed for 2 ns, followed by an additional 10,000 steps of minimization without any restraints. Further equilibration ensued for ∼4 ns with harmonic restraints on C_α_ atoms, followed by an additional unrestrained equilibration of ∼4 ns that allowed the complexes to equilibrate under the simulation conditions. Production MD simulations were run in the NPT ensemble using NAMD 3^50^ with the CHARMM36^51^ force field. Temperature was maintained at 310 K with a Langevin thermostat and pressure at 1 atm with a Langevin barostat, using a 2 fs time step. Long*-*range electrostatics were computed with the Particle Mesh Ewald (PME) method and a real-space cutoff of 12.0 Å for van der Waals interactions. Periodic boundary conditions were applied in all dimensions. Each RBD–Nb system was simulated in two independent 400-ns MD trajectories, yielding a total of 800 ns of simulation time per RBD–Nb complex and 10.4 μs across all complexes. Coordinates were sampled every 0.5 ns.

### PCA-based construction of binding-pose free-energy surfaces and basin-specific conformational ensembles

For each RBD–Nb complex, the two independent trajectories were concatenated, and all frames were aligned on the RBD C_α_ atoms to remove global translation/rotation and focus on relative interface motion. Principal component analysis (PCA)^22,23^ was then performed on the Nb C_α_ coordinates to quantify dominant variations in the Nb binding pose relative to the RBD. The covariance matrix^24^ was constructed using the aligned Nb coordinates:

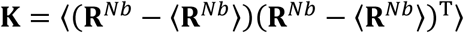

where **_R_***^Nb^* is the 3N-dimensional configurational vector composed of the instantaneous Nb C_α_ coordinates (N is the number of Nb residues) and 〈**_R_*^Nb^***〉 denotes the trajectory average. Eigenvalue decomposition of **_K_** yields principal components (PCs) **_p_**_i_ and their variances *_σ_*_i_:

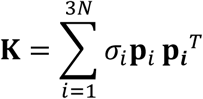

Each frame was projected onto the PCs as **_r*i*_** = **_p*i*_***^T^*(**_R_** − 〈**_R_**〉). We focused on PC1 and PC2, which capture the dominant Nb reorientation/translation relative to the aligned RBD (the sign of a PC axis is arbitrary). The joint probability density *_f_*(**_r1_**, **_r_**_2_) was converted to a two-dimensional potential of mean force (free-energy surface) using *_A_*(**_r1_**, **_r_**_2_) = −*_kTln_* (*_f_*(**_r1_**, **_r_**_2_)) + *_constant_*.^25^ Low-free-energy basins were defined by local minima in *_A_*(**_r1_**, **_r2_**) and the barriers separating them. Basin assignments were validated by monitoring CDR–RBD distances (**Figure S1**) to ensure that each basin corresponds to a distinct binding pose.

For complexes that sampled multiple binding poses, the trajectories were partitioned into basin-specific conformational ensembles corresponding to states S1–S3; for complexes with a single dominant basin S1, the full bound-state trajectory was retained as the analysis ensemble. Nb21 and Sb14 were excluded from the subsequent covariance-based interaction mapping because they dissociated from the RBD in both independent trajectories. The resulting bound-state ensembles for the remaining 11 complexes were used for the residue–residue interaction mapping described below.

### Covariance-based interaction mapping

We applied our covariance-matrix–based method^30,52^ to systematically map stabilizing (attractive) and destabilizing (repulsive) residue–residue interaction hotspots at the RBD–Nb interface from the MD simulations. The analysis proceeded in four steps:

*(i) Interprotein normalized covariance (cross-correlation matrix) calculation:* For each RBD–Nb complex, we computed the normalized covariance between the positional fluctuations of all Nb C_α_ atoms and all RBD C_α_ atoms. This yielded an N_Nb_ × N_RBD_ matrix (where N_Nb_ is ∼120 residues in the Nb and N_RBD_ is 195 residues in RBD), in which each element *C(i,j)* quantifies the correlation of motions between Nb residue *i* and RBD residue *j*

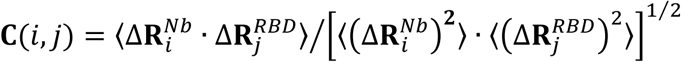

Here, 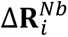 and 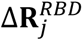 denote the displacement vectors of residues *i* and *j* (C_α_ atoms) from their time-averaged positions (after alignment), and ⟨·⟩ denotes a time average over the trajectory. Positive **_C_**(*_i_*, *_j_*) indicates correlated motion, whereas negative **_C_**(*_i_*, *_j_*) indicates anticorrelated motion. For residue pairs that remain spatially proximal at the interface, strong positive or negative correlations can serve as signatures of persistent attractive interactions or dynamically coupled repulsive interactions, respectively. Residue pairs with large |**_C_**(*_i_*, *_j_*)| values are prioritized as candidate hotspots.
*(ii) Spatial filtering to generate the close-contact covariance matrix:* Because correlations can also arise from indirect/long-range coupling, we filtered candidate residue pairs by proximity. We retained only RBD–Nb residue pairs whose mean C_α_-C_α_ distance over the simulation was ≤11 Å for positively correlated pairs and ≤13 Å for negatively correlated pairs; all other pairs were discarded, yielding a close-contact covariance matrix.
*(iii) Interaction identification and classification:* For residue pairs retained in the close-contact covariance matrix, we annotated the underlying interaction type(s) using geometric criteria evaluated across MD conformations. Attractive interactions were classified as salt bridges (oppositely charged side-chain groups within ∼6 Å),^53^ hydrogen bonds (donor–acceptor distance ≤3.5 Å and donor–H–acceptor angle ≥150°),^54^ and hydrophobic interactions (nonpolar side chains within ∼8 Å).^55^ Repulsive interactions were flagged as same-charge electrostatic repulsion (like-charged side-chain groups within 12 Å) and hydrophobic–charged polarity mismatch (an aliphatic side chain within ∼12 Å of a charged group).
*(iv) Mapping interaction frequencies:* For each classified residue pair, we computed its occupancy (percentage of frames satisfying the corresponding geometric criterion), yielding an interaction-frequency map for each RBD–Nb complex.

All steps were implemented using custom MATLAB scripts; VMD^56^ was used for trajectory processing and visualization.

### Steered molecular dynamics simulations of force-induced unbinding

To estimate and compare the mechanical resistance of ACE2 and Nbs to forced detachment from the Omicron RBD, we performed low-speed SMD unbinding simulations as an *in silico* analogue of single-molecule force spectroscopy. In this approach, a dummy atom is moved with constant velocity along a predefined pulling direction, and the steered atoms are coupled to this dummy atom through a virtual spring. The instantaneous force and the accumulated mechanical work required for unbinding provide a quantitative measure of force-induced detachment and enable relative comparisons among binders under identical simulation settings.

All SMD simulations were performed at a constant pulling velocity of 0.1 Å ns⁻¹ with a spring constant of 100 kcal mol⁻¹ Å⁻², consistent with a stiff-spring regime in which the center of mass of the steered group follows the dummy atom closely while still allowing thermal fluctuations. The guiding potential is defined as

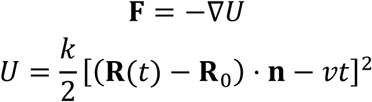

where *_U_* is the harmonic guiding potential, *_k_* is the spring constant, *_v_* is the pulling velocity, *_t_* is time, **_n_** is the unit vector defining the pulling direction, **_R_**(*_t_*) is the instantaneous center-of-mass position of the steered atoms, and **_R_**_0_ is their initial center-of-mass position. The mechanical work along the reaction coordinate *_ξ_* was computed by integrating the applied force along the pulling displacement,

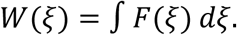

During the SMD simulations, C_α_ atoms were used to define both steered and fixed atom groups. The steered atoms were defined as all C_α_ atoms of ACE2 for RBD–ACE2 pulling, and all C_α_ atoms of the Nb for RBD–Nb pulling. The fixed atoms were defined as C_α_ atoms of three Omicron RBD segments (Y396–I402, C432–W436, and R509–F515), which served as an internal structural anchor for the RBD throughout pulling. The pulling direction **_n_** was defined as the vector from the center of mass of the fixed atoms (RBD anchor group) to the center of mass of the steered atoms (ACE2 or Nb) at the start of each simulation.

SMD simulations were performed for ACE2 and 11 Nbs: H11-D4, H11-H4, H3, MR17, NM1230, Re5D06, Sb23, Sb45, Ty1, VHH-E, and WNb-2. Each SMD simulation was run for 300 ns, corresponding to an overall pulling distance of approximately 30 Å. Four independent SMD trajectories were initiated from four distinct bound conformations for each of the 11 RBD–Nb complexes and also for the RBD-ACE2 complex. For each RBD–Nb complex, the starting conformations were sampled from the two equilibrium conventional MD trajectories generated in this study, whereas the ACE2–RBD starting conformations were sampled from the equilibrium MD simulations of the Omicron BA.2 RBD–ACE2 complex reported previously.^8^ The same SMD protocol described above was applied to all systems.

Attractive interactions along the SMD unbinding pathways were identified using the geometric criteria described above. Because interactions formed during force-induced unbinding were often transient, attractive interactions present in ≥1% of frames within each complete SMD simulations were retained. In contrast, the conventional-MD bound-state analysis used reporting thresholds of ≥49% for attractive interactions. Intermediate bound conformations were selected from individual SMD trajectories as structurally distinct, partially bound poses associated with local force peaks followed by force drops. For each selected snapshot, attractive residue-pair interactions were classified as retained, lost, or newly formed relative to the initial bound state and, where indicated, the preceding intermediate.

### RMSD statistical analysis

After alignment of each frame on the RBD C_α_ atoms, the Nb C_α_ RMSD was calculated relative to the corresponding experimentally determined reference pose for each of the 13 systems. To reduce the influence of temporal autocorrelation, each RMSD trajectory was divided into non-overlapping blocks of 100 frames, corresponding to 50 ns, and the mean RMSD was calculated for each block. This yielded 16 block-averaged RMSD values per system, with eight values from each independent 400-ns trajectory. The resulting block-averaged RMSD values for each pose-plastic Nb (Nb21 and Sb14) were compared separately with those of each of the 11 single-basin Nbs using two-sided Welch’s independent-samples t-tests.

## RESULTS AND DISCUSSION

### Molecular dynamics simulations reveal Omicron-dependent shifts in nanobody binding-pose free-energy landscapes

To probe how Omicron-associated mutations in the SARS-CoV-2 spike RBD affect Nb binding poses, we performed all-atom MD simulations for 13 RBD–Nb complexes: H11-D4, H11-H4, H3, MR17, Nb21, NM1230, Re5D06, Sb14, Sb23, Sb45, Ty1, VHH-E, and WNb-2 (**Table 1**). Starting from experimentally determined Nb–RBD structures,^5,38–47^ we introduced the Omicron BA.2 mutations *in silico* and carried out two independent 400-ns simulations for each complex (800 ns per Nb). The RBD sequence was kept fixed as the Omicron BA.2 RBD in all complexes; therefore, the variable component across the panel was the Nb sequence and binding geometry. The Nbs differ in CDR sequences, length, and chemistry, especially in CDR3 (**Figure 1A**), as well as in their framework regions.

MD simulation trajectory frames were aligned by least-squares fitting of the RBD C_α_ atoms to the WT RBD reference structure, allowing us to focus on Nb reorientation relative to the spike RBD. We then quantified these pose changes using PCA of the Nb C_α_ coordinates from the aligned trajectories and projected all sampled conformations onto the first two principal components (PC1 and PC2). The resulting conformational densities were converted to two-dimensional free-energy landscapes (**Figure 2**).

**Figure 2.**
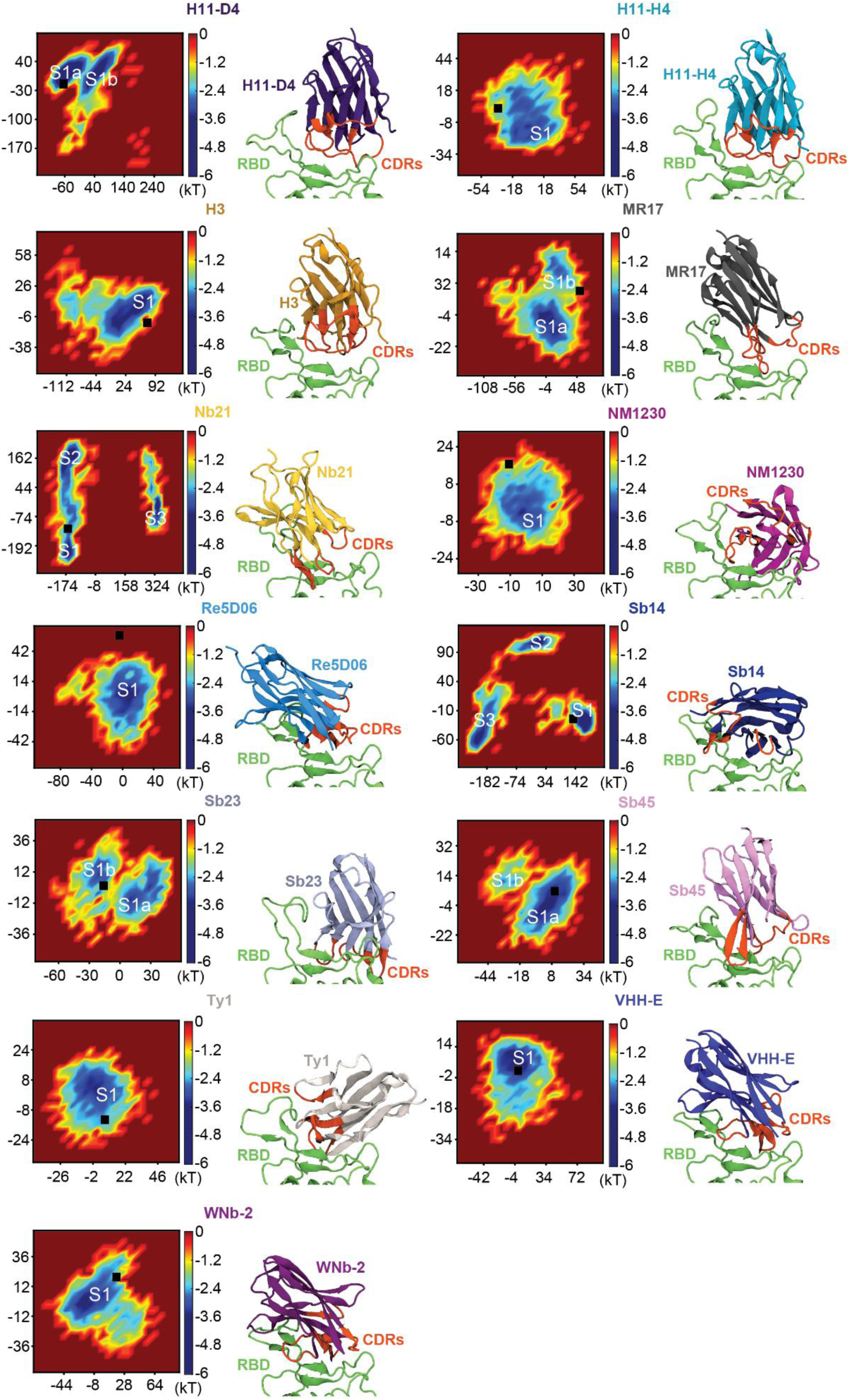
Nbs sample single or multiple binding poses on the Omicron SARS-CoV-2 RBD. For each RBD–Nb complex, two-dimensional free-energy surfaces are shown in the space of PC1 and PC2, describing binding-pose variability. Free energies are plotted in units of kT and shifted by an arbitrary additive constant; the color scale ranges from 0 (red) to −6 kT (dark blue), with more negative values indicating more highly populated states. Dominant low-free-energy basins are labeled S1–S3 and used to define the most populated binding-pose states. In each panel, the black square indicates the experimentally determined reference (all WT RBD, except Alpha/Kent RBD in Nb H3) binding pose projected into the corresponding PC1–PC2 space. The position of the reference-pose marker relative to the dominant Omicron basin provides a qualitative measure of pose preservation in this reduced coordinate system: proximity to the basin center indicates strong preservation, placement near the basin boundary indicates partial adjustment, and separation from the populated basin region indicates substantial rearrangement. Representative conformations taken from the free-energy minimum (darkest blue region) of the S1 basin are shown to the right of each free-energy surface (RBD, green; Nb, shown in distinct colors; CDR loops, orange/red).

Across the Nb panel, 11 Nbs (H11-D4, H11-H4, H3, MR17, NM1230, Re5D06, Sb23, Sb45, Ty1, VHH-E, and WNb-2) remained largely confined to a single dominant low free energy basin (S1), corresponding to one dominant binding state, with varying degrees of within-state heterogeneity and intra-state substates. Four of these 11 Nbs (H11-D4, MR17, Sb23, and Sb45) showed two major binding substates, S1a and S1b, which exhibited slightly different Nb orientations (**Figure S2**) while preserving a similar RBD-CDR interfacial distance (**Figure S1**). For the other Nbs, additional substates were sampled much less frequently than the main state and therefore were not separately annotated. By comparison, Nb21 and Sb14 populated three well-separated basins (S1– S3) and dissociated from the RBD in both independent trajectories (**Figure S2**), demonstrating pronounced pose plasticity and the absence of a persistent bound state under Omicron.

The extent to which the investigated Nbs preserved their experimentally determined reference poses after introduction of the Omicron BA.2 substitutions varied substantially (**Figure 2** and **Figure S2**). Except for H3, all experimentally determined reference structures contained the ancestral WT RBD. The H3 reference structure contained the Alpha/Kent RBD carrying the N501Y substitution; however, residue 501 does not directly contact H3, and the original structural study found no detectable N501Y-induced change in the RBD-H3 interface.^38^ Among the Nbs that retained a single dominant binding state, the corresponding reference poses projected within the dominant Omicron basin for only five binders: H11-D4 and Sb45 within S1a, and VHH-E, Ty1, and WNb-2 within S1. For H11-H4, H3, and NM1230, the corresponding reference poses projected near the outer boundary of S1; for MR17, its reference pose projected near the outer boundary of S1b, whereas for Sb23, the corresponding reference pose projected near the barrier separating S1a and S1b, but closer to S1b. In contrast, the reference pose for Re5D06 projected outside the most populated Omicron region. For the multi-basin cases, the reference pose projected within the S1 basin, corresponding to the first binding state sampled in these simulations.

The mean C_α_ RMSD of Omicron-sampled Nb poses relative to the corresponding experimentally determined reference poses ranged from 2.5-8.2 Å for single-basin Nbs (Table S1), whereas the pose-plastic binders showed greater variability and significantly larger mean RMSDs: Sb14 (18.6 ± 10.7 Å; p values ranging from 2×10^-5^ to 10^-3^ across pairwise comparisons with the single-basin Nb RMSD distributions) and Nb21 (27.9 ± 15.5 Å; p values ranging from 7×10^-6^ to 1×10^-4^ across the corresponding comparisons). Although methodological and thermodynamic differences between experimentally determined reference structures and MD simulations performed under physiologically relevant solution conditions may contribute to the absolute displacements between the simulated Omicron ensembles and their starting reference structures,^57^ the broad Nb-dependent variation in reference-pose projection and RMSD supports the conclusion that Omicron mutations alter the binding-pose ensembles of several Nbs.

### Covariance fingerprints reveal Nb binding footprints and stabilizing and destabilizing interfacial hotspots

Because Nb21 and Sb14 dissociated from the RBD in both independent trajectories, the covariance analysis was restricted to the 11 complexes that retained a persistent bound state. For these complexes, the full bound-state trajectory was used as the analysis ensemble. Interpreting multi- µs long MD trajectories of RBD–Nb complexes poses a major data-reduction challenge, because hundreds of residue pairs can contribute to binding, making it difficult to distinguish persistent, dynamically meaningful interactions from incidental geometric contacts. To enable a systematic and scalable comparison across the Nb panel examined here, we employed our recently developed covariance-matrix-based interaction-mapping method.^30,52^ This approach constructs an interprotein normalized covariance (C_α_-C_α_ cross-correlation) matrix from the aligned trajectories, applies a distance-based filter to focus on physically plausible contacts, and then identifies the attractive (stabilizing) and repulsive (destabilizing) interactions underlying these correlations using the geometric criteria described in Methods. By prioritizing dynamically coupled residue pairs that remain spatially proximal, the workflow efficiently pinpoints the residue–residue interactions that stabilize or destabilize Nb binding. The resulting close-contact covariance matrices provide compact, dynamic “at-a-glance” fingerprints of each RBD–Nb interface, showing where each Nb couples to the RBD surface and whether the associated interfacial motions are correlated or anticorrelated (**Figure 3**).

**Figure 3.**
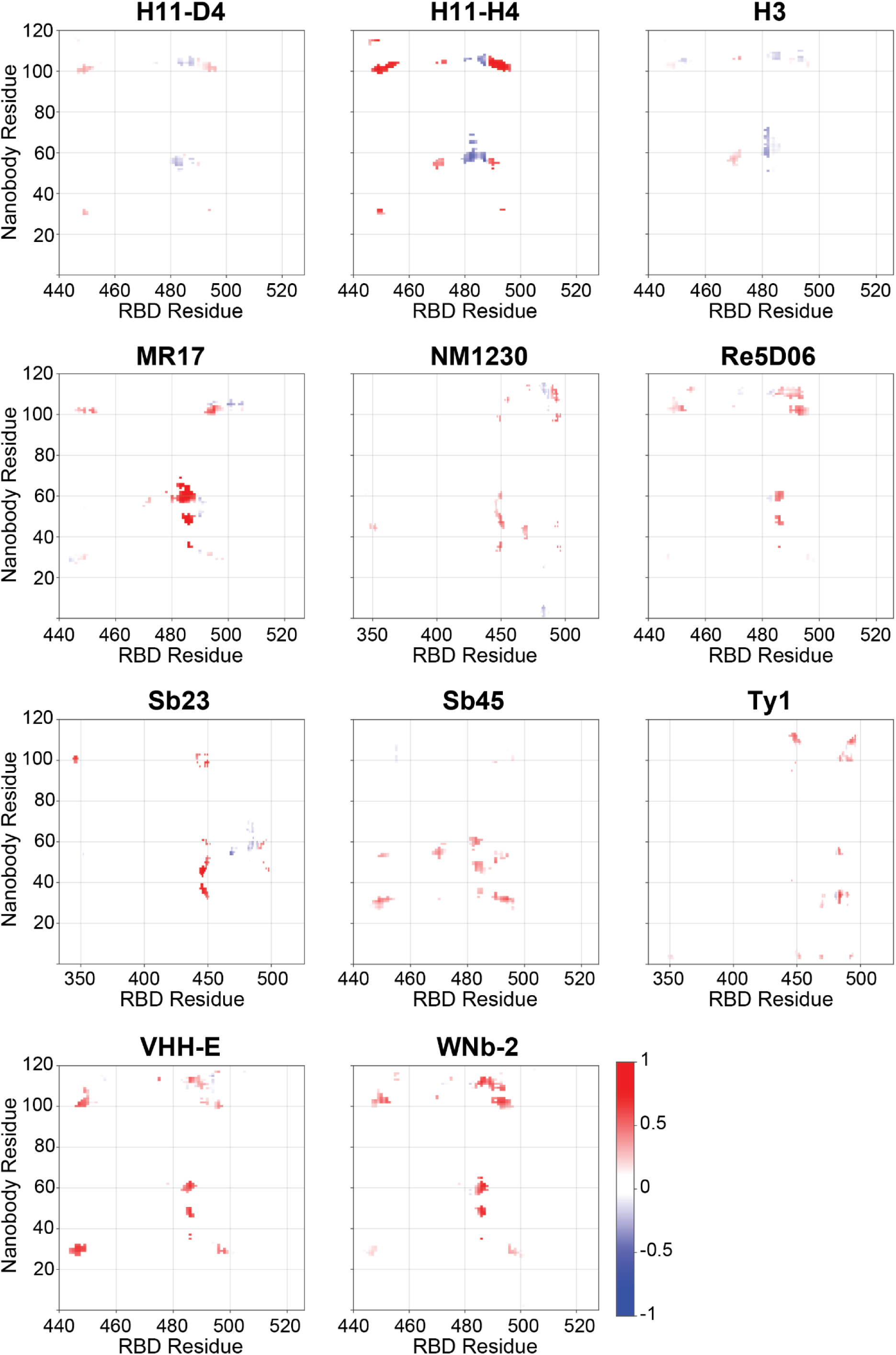
Interprotein cross-correlation for Omicron RBD–Nb complexes. Each panel is a close-contact covariance matrix **_C_** mapping residue–residue correlations between the Omicron RBD (x-axis, residue index) and the Nb (y-axis, residue index). In these two-dimensional maps, red regions correspond to residue pairs whose movements are positively correlated, indicating correlated interfacial motions (**_C_**(_i_, _j_) > 0), whereas blue regions highlight residue pairs that move in opposite directions and exhibit anticorrelated motions (**_C_**(_i_, _j_) < 0). Only residue pairs that come within contact range are shown (distant pairs filtered out), yielding close-contact covariance matrices that highlight dynamically coupled interfacial hotspots that were subsequently annotated as stabilizing or destabilizing based on interaction geometry.

These fingerprints reveal two coarse footprint classes across the Nbs analyzed here: most binders (H11-D4, H11-H4, H3, MR17, Re5D06, Sb45, VHH-E, and WNb-2) show RBM-focused coupling confined to the ACE2-contacting surface (∼S438–Q506), whereas a smaller subset (NM1230, Sb23, and Ty1) retains RBM anchoring but additionally engages N-terminal/core residues (∼E340– R355), consistent with broader binding footprints. Within the RBM focused class, covariance hotspots further allow subdivision by RBM sub-element: S443-N450 loop-engaging binders (VHH-E and WNb-2), a flat surface–biased binder (Re5D06) whose dominant contacts track the intervening planar RBM face, and ridge-weighted binders (H11-H4, H3, MR17, and Sb45) with substantial coupling across the receptor binding ridge (RBR; T470–C488). These footprint classes are defined by the spatial distribution of covariance hotspots, whereas the net covariance score (S_cov_), defined as the algebraic sum of all elements in the spatially filtered close-contact covariance matrix, provides a complementary quantitative summary of the overall balance between correlated and anticorrelated interfacial coupling. The S_cov_ scores showed similar trends across independent trajectories (**Figure S3; Table S2**) while remaining sensitive to trajectory-specific events. H11-D4 showed a larger S_cov_ difference between its two independent trajectories because of a transient unbinding event in one trajectory (**Figure S1**). WNb-2 and MR17 showed strongly positive S_cov_ values, while H3 had the lowest S_cov_, which was negative, consistent with a comparatively larger contribution from anticorrelated coupling and a low level of correlated coupling (**Figure 3**). This interpretation is also consistent with the interaction-frequency analysis (**Figure 4**; **Table 2**), in which H3 shows the largest repulsive-contact burden under the reporting thresholds.

**Figure 4.**
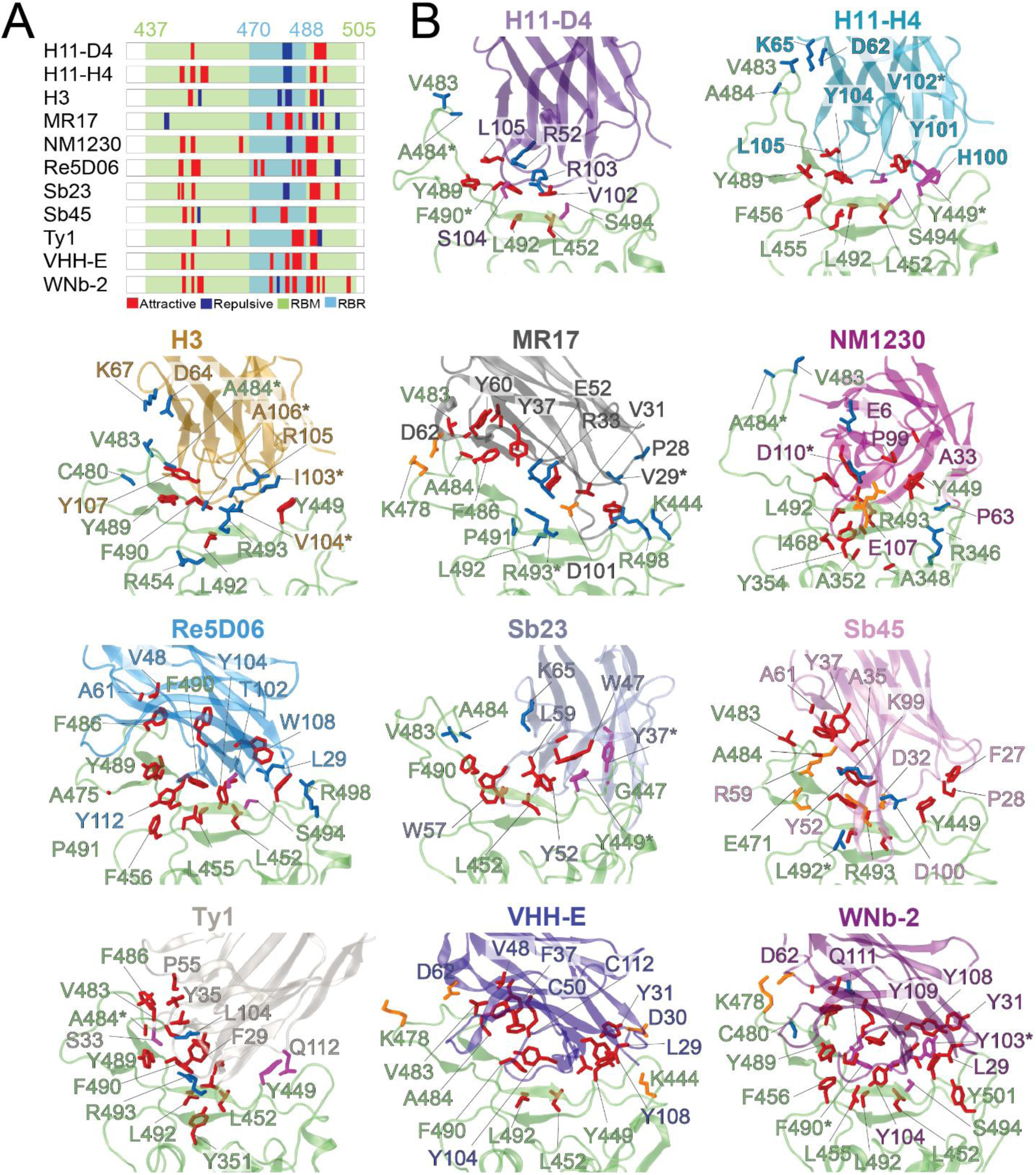
Structural mapping of Omicron RBD–Nb interfaces within the RBM. **(A)** Linear map of residue-level interaction profiles for RBD-binding Nbs. The SARS-CoV-2 spike sequence from N437 to H505 encompasses the RBM, shown in green, with the receptor-binding ridge (RBR; T470–C488) highlighted in light blue. Colored bars indicate residues contributing attractive (red) or repulsive (blue) interactions between each Nb and the RBD. **(B)** Structural visualization of Nbs bound to the Omicron RBD (green). Interacting RBD residues are colored by interaction type: hydrophobic interaction (red), salt bridge (orange), hydrogen bond (magenta), and repulsive interaction (blue). Only repulsive interactions with ≥14% occupancy and attractive interactions with ≥49% occupancy over the full simulation are shown. Asterisks indicate residues involved in multiple electrostatic interactions (**Table S3**).

**Table 2.** Interactions between Nbs and the SARS-CoV-2 Omicron Variant RBD. Grouped interactions for the Nbs considered here based on high-frequency (>49%) attractive interactions (salt bridge, SB; hydrogen bond, HB; hydrophobic, HP) and medium-to-high (>14%) repulsive interactions (R; like-charge electrostatic repulsion and hydrophobic–charged polarity mismatch).

| Nanobody | SB | HB | HP | R |
| --- | --- | --- | --- | --- |
| **H11-D4** | 0 | 2 | 4 | 3 |
| **H11-H4** | 0 | 2 | 8 | 4 |
| **H3** | 0 | 0 | 7 | 12 |
| **MR17** | 2 | 0 | 9 | 6 |
| **NM1230** | 2 | 0 | 12 | 3 |
| **Re5D06** | 0 | 1 | 14 | 1 |
| **Sb23** | 0 | 1 | 11 | 2 |
| **Sb45** | 2 | 0 | 12 | 2 |
| **Ty1** | 0 | 2 | 15 | 1 |
| **VHH-E** | 1 | 0 | 13 | 0 |
| **WNb-2** | 1 | 3 | 18 | 1 |

Interaction-frequency mapping (**Figure 4**; **Table 2**; **Figures S4** and **S5**) shows that, across Nbs, stabilizing interactions are overwhelmingly hydrophobic and converge on a recurrent RBM “anchor” centered near V445–F456 and F490–Y501, with variable reinforcement from hydrogen bonds or salt bridges. Stabilizing interactions that engage Omicron-substituted sites include T478K (MR17, VHH-E, WNb-2), E484A (Sb45, Ty1, VHH-E), Q493R (MR17, NM1230, Sb45), and N501Y (WNb-2). Conversely, Omicron-substituted sites also participate in destabilizing repulsive interactions at E484A (H11-D4, H11-H4, H3, NM1230, Sb23), Q493R (H3, MR17, Ty1), and Q498R (Re5D06). The main difference among Nbs is the extent to which the shared RBM anchoring is offset by RBR-localized (and adjacent) repulsive interactions around ∼Q474–A484 (often involving E484A): Re5D06, VHH-E, WNb-2, and Ty1 are nearly repulsion-free above the reporting threshold, whereas H3 exhibits persistent polarity/charge mismatches that erode the net coupling, with similar mismatch features also observed (to a lesser extent) for H11-H4, MR17, Sb45, NM1230, and Sb23. Overall, covariance fingerprints connect epitope footprint (RBM-only vs core-spanning) to binding robustness (hydrophobic anchoring with minimal repulsion versus mutation-adjacent repulsion) in a single view. Residue-level interaction partners, occupancies, and mutation-specific interactions for each Nb are provided in the **supplementary materials**.

### Nb resistance to unbinding from the Omicron RBD is substantially lower than that of ACE2

To directly compare the relative stability of ACE2 and Nb binding to the Omicron RBD, and to characterize the unbinding process in terms of interaction breakage and formation, intermediate binding poses, binding anchors, and resistance to unbinding, we performed low-speed SMD unbinding simulations of ACE2 and the 11 Nbs that retained persistent bound states during conventional MD. In this analysis, resistance to unbinding was quantified as the average total external work required to detach each binder under the same pulling protocol. We applied a pulling speed of 0.1 Å ns⁻¹, which is within the range used in high-speed AFM-based force spectroscopy experiments (**Figure 5**). SMD unbinding trajectories are provided as examples in **Videos S1–S12** and were rendered with UCSF ChimeraX^58^. Among the systems tested, ACE2 exhibited the highest resistance to detachment, with its work curve lying above the Nb curves throughout the reaction coordinate (**Figure 5B**), indicating that ACE2 forms the strongest complex with the Omicron RBD. The average work required to unbind ACE2 from the Omicron RBD is 45 kcal mol⁻¹ (**Table 3**). In contrast, all tested Nbs required less mean work to detach from the Omicron RBD (16-42 kcal mol⁻¹). For all but two Nbs, the difference was substantial, amounting to approximately 10-30 kcal mol⁻¹ less work than for ACE2. These results indicate that the tested RBD–Nb interfaces are mechanically easier to disrupt under force than the RBD–ACE2 interface. In particular, Nbs H11- D4, H11-H4, and Ty1, which were previously shown^4^ to exhibit work values comparable to or slightly higher than those for ACE2 on the WT RBD, detach from the Omicron RBD with 15.5, 22.7 and 25.8 kcal mol⁻¹, respectively (**Table 3**), corresponding to ∼34.5%, ∼50.4%, and ∼57.2% of the ACE2 work value. Other Nbs quantified here also had lower mean work values than ACE2, including H3 (23.8 kcal mol⁻¹), VHH-E (29.4 kcal mol⁻¹), and Sb45 (31.0 kcal mol⁻¹), with H11- D4 (15.5 kcal mol⁻¹) exhibiting the lowest mean unbinding work. Sb23 (41.8 kcal mol⁻¹) and NM1230 (42.4 kcal mol⁻¹) approached the ACE2 work value most closely, although both remained lower (**Table 3**).

**Figure 5.**
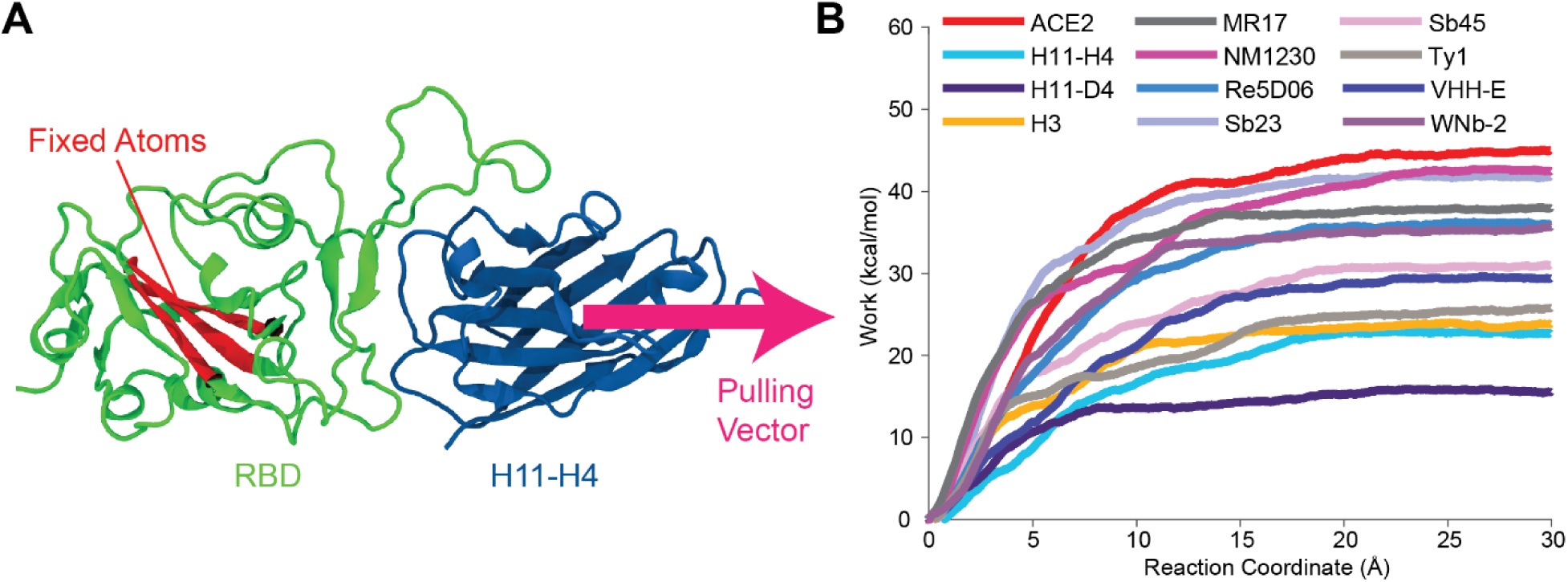
Force-induced detachment of ACE2 and Nbs from the Omicron RBD. **(A)** Schematic of the SMD pulling setup, shown for the RBD–H11-H4 complex as a representative example. The fixed atoms (red) are the RBD C_α_ atoms of residues Y396–I402, C432–W436, and R509–F515. The steered atoms consist of all C_α_ atoms of the bound partner (ACE2 or Nb), and the pulling direction (magenta arrow) is defined by the vector from the center of mass of the fixed group to the center of mass of the steered group at the start of each simulation. **(B)** Average accumulated unbinding work, _W_(_ξ_), as a function of the reaction coordinate _ξ_ (center-of-mass displacement of the steered group along the pulling direction) obtained from SMD simulations performed at a constant pulling velocity of 0.1 Å ns^-1^ (spring constant _k_ = 100 kcal mol^-1^ Å^-2^). Final work values (mean ± s.d. over four independent trajectories per system) are summarized in **Table 3**.

**Table 3.** Unbinding work values of the RBD-ACE2 and the RBD–Nb systems.

| System | Work (kcal mol<sup>-1</sup>), mean ± s.d. | Work (kT), mean ± s.d. |
| --- | --- | --- |
| ACE2 - RBD | 45.0 ± 11.7 | 73.0 ± 19.0 |
| H11-D4 - RBD | 15.5 ± 9.1 | 25.2 ± 14.8 |
| H11-H4 - RBD | 22.7 ± 6.4 | 36.8 ± 10.3 |
| H3 - RBD | 23.8 ± 7.2 | 38.7 ± 11.7 |
| MR17 – RBD | 38.0 ± 3.6 | 61.6 ± 5.9 |
| NM1230 – RBD | 42.4 ± 7.2 | 68.9 ± 11.7 |
| Re5D06 – RBD | 36.0 ± 4.4 | 58.5 ± 7.1 |
| Sb23 – RBD | 41.8 ± 11.6 | 67.9 ± 18.9 |
| Sb45 – RBD | 31.0 ± 5.4 | 50.3 ± 8.8 |
| Ty1 – RBD | 25.8 ± 8.1 | 41.8 ± 13.2 |
| VHH-E – RBD | 29.4 ± 10.2 | 47.7 ± 16.6 |
| WNb-2 – RBD | 35.7 ± 14.1 | 57.9 ± 22.9 |

Experimental Omicron benchmarking is available for only a small subset of the Nbs investigated here; however, directly comparable affinity measurements for the original monomeric Nbs on BA.2 are not available. Instead, the existing evidence consists mainly of neutralization/escape data and some relative binding assays for engineered constructs or mutation panels.^40,59,60^ Direct neutralization data exist for Nb21-Fc, which neutralizes WT and Delta but fails against authentic BA.1 and pseudotyped BA.1/BA.1.1/BA.2, whereas Ty1-, H11-H4-, and MR17-K99Y-derived modules have been evaluated mainly as components of engineered multimodular constructs against BA.1, with markedly reduced neutralization compared with WT.^40,59,60^ Furthermore, these studies do not provide a matched head-to-head experimental comparison of the original monomeric Nbs with ACE2 on the same BA.2 RBD. Thus, while our combination of conventional and steered MD simulations are qualitatively consistent with the limited Omicron benchmarks available for Nb21-Fc and for Ty1-, H11-H4-, and MR17-K99Y-derived modules, they also extend the analysis to Nbs lacking comparable Omicron data. The SMD analysis further fills a critical gap by providing a uniform ACE2-versus-Nb comparison for the 11 BA.2 RBD–Nb systems that retained persistent bound states during conventional MD simulations.

### Pathway-resolved SMD reveals stepwise unbinding through transiently anchored intermediates

In addition to directly comparing the resistance of ACE2 and the Nbs to force-induced unbinding from the Omicron RBD, our SMD simulations provided mechanistic insight into the unbinding process itself. Unbinding did not occur as a single catastrophic rupture event. Instead, detachment proceeded through a sequence of interaction-breakage and reorganization events during which specific interfacial patches remained engaged after other interactions had been lost. These persistent patches generally involved hydrophobic interactions, which acted as transient anchors during dissociation. Their delayed rupture generated short-lived, partially bound intermediates that increased the mechanical work required for complete detachment, thereby raising the effective energy barrier encountered during unbinding.

Across the SMD unbinding trajectories, individual RBD–Nb complexes followed heterogeneous pathways and sampled various intermediates. Nevertheless, RBR-mediated anchoring emerged as a recurring feature that stabilized partially bound conformations and maintained Nb attachment to the RBD. Within this recurring pattern, the RBD C480-F486 segment remained engaged with the CDR2 and CDR3 regions of most Nbs until the late stages of unbinding (**Figures 6** and **S6** and **Videos S2-S12**). For example, in one H11-H4 trajectory, persistent interactions with the S443- N450 loop and the flat-surface regions N450-F456 and C488-F497 were lost at a pulling distance of approximately 11 Å, after which the Nb slightly rearranged its binding mode and engaged the RBR through the I472-K478 residue stretch. These interactions persisted until a pulling distance of approximately 17.5 Å, forming a transient intermediate stabilized by hydrogen bonds and hydrophobic interactions. This re-engagement coincided with a force peak and a corresponding rise in accumulated work. Similarly, in one Re5D06 trajectory, rupture of the primary interface was followed by partial rebinding as the Nb moved along the A484-F486 region of the RBD, sequentially breaking existing interfacial interactions and forming new ones. This interaction exchange coincided with the late-stage rise observed in the Re5D06 work profile. In some Nb unbinding trajectories, including all of the H3 trajectories, the RBR anchoring by CDR2 and CDR3 was further assisted by the Nb’s framework residues adjacent to CDR2. Together, these observations identify RBR-mediated anchoring as a recurring mechanism by which Nbs maintain partial attachment while transiently forming new interactions during force-induced unbinding despite differences in their initial binding poses.

**Figure 6.**
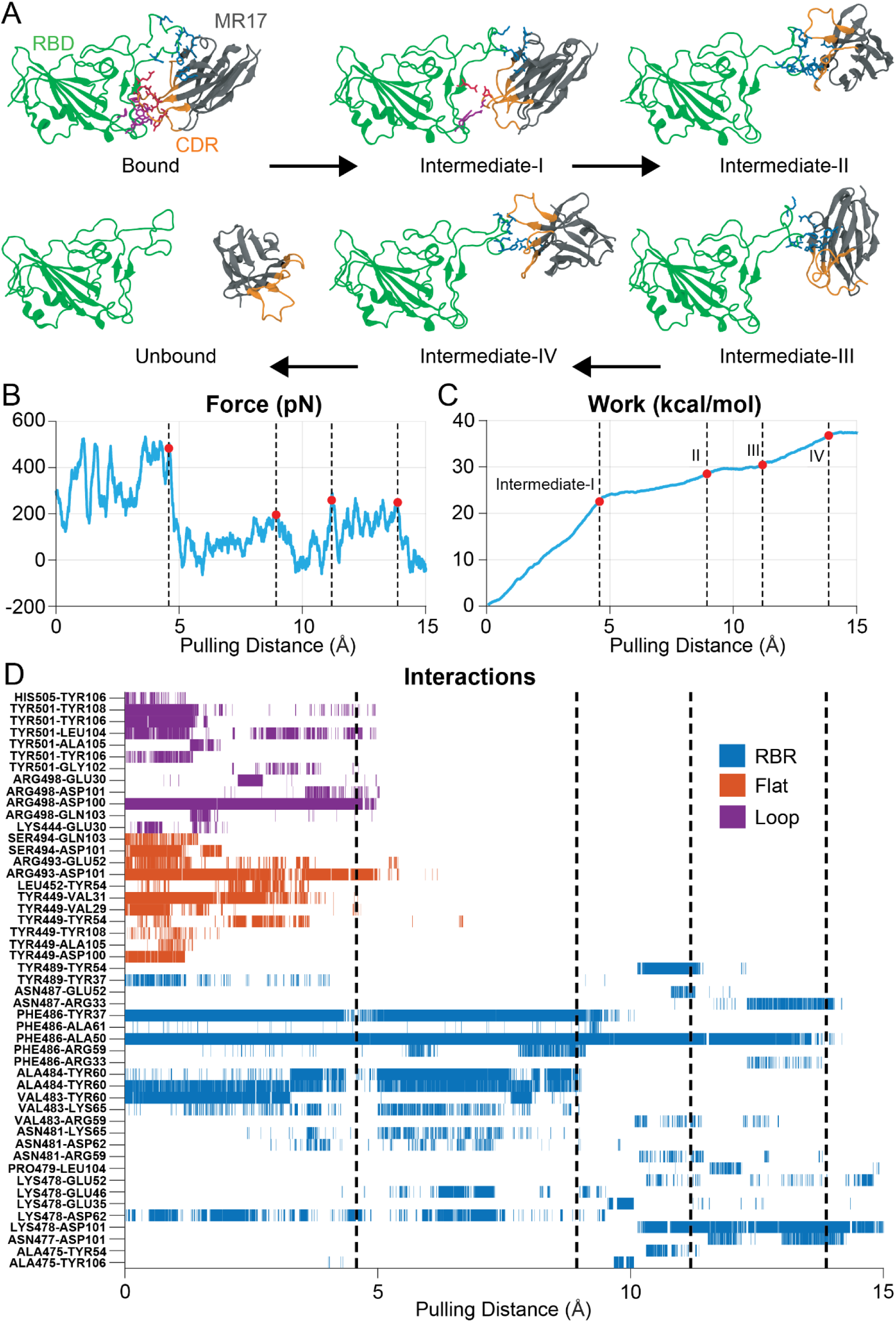
Stepwise unbinding of MR17 from the Omicron RBD through transient intermediates. (A) Structural snapshots from one SMD trajectory showing the bound state, four intermediate conformations (I–IV), and complete dissociation. The RBD is shown in green and MR17 in gray, with the CDR loops highlighted in orange. RBD residues remaining at the interface are shown in licorice representation and colored by region: the receptor-binding ridge (RBR), blue; flat surface, red; and S443–N450 loop, purple. Arrows indicate progression along the pulling pathway. **(B)** Force, **(C)** accumulated work, and **(D)** attractive interactions as functions of pulling distance for the same trajectory. Red circles mark the intermediate conformations shown in panel A. Local force peaks and subsequent drops accompany successive interaction rearrangement and rupture events, whereas accumulated work increases stepwise and plateaus after complete separation.

To illustrate the stepwise nature of unbinding at residue-level resolution, we present one MR17 unbinding trajectory (**Figure 6**). The structurally distinct, partially bound intermediates overlapped with changes in the force profile, including local force peaks followed by force drops as the anchoring interactions stabilizing each pose ruptured. The initial bound state contained 19 attractive interactions distributed across the RBR, flat-surface and loop regions of RBD. At a pulling distance of approximately 5 Å, MR17 reoriented to form intermediate I, losing 11 attractive interactions present in the initial bound state while forming two new attractive interactions, resulting in a reorganized network of ten interactions (**Figures 6** and **S6**). Further pulling from 5 to 9 Å disrupted most of the remaining interactions inherited from the initial bound state, including all interactions with the Flat and Loop regions, while newly formed interactions stabilized intermediate II (**Figures 6** **and S6**). Between 9 and 12 Å, further interaction breakage and binding-pose rearrangement, accompanied by the formation of seven new RBR interactions, generated intermediate III. At later pulling distances, MR17 sampled intermediate IV at approximately 14 Å before complete dissociation. Although the specific interaction changes and MR17 reorientations differed among transitions, repeated engagement of the RBR by MR17’s all three CDR loops was the recurring anchoring feature. Thus, this trajectory illustrates how RBR-mediated anchoring stabilizes successive intermediates that delay complete force-induced unbinding.

## CONCLUSION

In this study, we combined all-atom MD simulations with a covariance-matrix–based interaction mapping framework to dissect how Omicron (BA.2) RBD substitutions reshape Nb recognition at atomic resolution. Analysis of PCA-derived binding-pose free-energy surfaces revealed a spectrum of Nb binding-pose behavior under Omicron. Eleven Nbs retained a dominant bound basin but showed varying degrees of within-basin heterogeneity and also differed in how closely their Omicron-bound poses matched the corresponding experimentally determined reference poses, which, except for H3, were obtained from structures containing the ancestral WT RBD. In contrast, Nb21 and Sb14 exhibited pronounced pose plasticity, populated multiple well-separated basins, and dissociated from the RBD, indicating loss of a persistent bound state. These results emphasize that Omicron can weaken Nb binding not only through the loss of individual interactions but also through mutation-driven reorientation and increased heterogeneity of the bound ensemble.

Our covariance-based fingerprints and interaction-frequency maps linked these pose-level changes to residue-level mechanisms. Across the 11 complexes that retained persistent bound states, stabilizing interactions were dominated by hydrophobic interactions that converged on recurrent RBM anchor patches near V445-F456 and F490-Y501, with binder-specific reinforcement from hydrogen bonds and salt bridges. Many of these recurrent interaction networks were rewired by Omicron substitutions and associated local structural rearrangements (e.g., in regions surrounding residues 478, 484, 493, 498, and 501), producing distinct balances of stabilization and destabilization depending on CDR chemistry and local complementarity. Nbs differed most strongly in the extent to which repulsive interactions offset shared hydrophobic anchoring. This repulsive burden was minimal for Re5D06, VHH-E, WNb-2, and Ty1 but more pronounced in the remaining Nbs, which exhibited persistent polarity mismatches and/or like-charge repulsion, particularly near the ridge-adjacent Q474–A484 region containing the E484A site. Thus, Omicron-driven weakening can frequently be explained by the introduction of unfavorable interactions and anticorrelated interfacial motions, rather than simply by the loss of favorable interactions.

The SMD provided a complementary measure of mechanical robustness that aligned with these interaction-network trends to mechanical robustness. Unlike endpoint structure-based energy estimates, the unbinding work values were obtained directly from the external bias applied during the simulations by recording the steering force and integrating it along the pulling coordinate; therefore, they are trajectory-defined mechanical observables rather than post hoc predictions from selected conformations. The molecular observations during the unbinding process highlighted the added value of SMD beyond end-state structural or energetic analyses. Endpoint-based approaches can identify the interactions present in the fully bound complex, but they do not reveal the order in which these interactions rupture, whether partially bound intermediates are formed, which patches resist detachment the longest, or how local rearrangements contribute to the total unbinding work. In contrast, the SMD trajectories resolve the unbinding pathway directly and show that the mechanical stability of each complex is determined not only by the number of interactions in the initial bound state, but also by the persistence, rearrangement, and sequential rupture of anchor-like interfacial patches during dissociation.

At the pathway level, the SMD trajectories identified RBR-mediated anchoring as a recurring mechanism that stabilized partially bound intermediates and delayed complete force-induced unbinding. At the system level, ACE2 required the largest mean work to unbind from RBD, whereas all tested Nbs detached with lower mean work values. Within the Nb set, NM1230 and Sb23 exhibited the highest mean resistance to detachment, whereas H11-H4, H11-D4, and Ty1 (whose WT work values were previously comparable to or slightly higher than those for ACE2) showed markedly lower work values on Omicron, indicating that Omicron shifts the force-resistance balance in favor of ACE2. The overall work trends in overall aligned with the covariance-guided interaction analyses: Nbs with extensive, persistent attractive networks and minimal repulsion exhibited greater mechanical resistance to unbinding, whereas mutation-adjacent repulsive features were associated with easier disruption of the interface. Because ACE2 binds the RBD with high affinity, Nbs that neutralize the virus by directly competing with or sterically blocking the ACE2-binding site must effectively outcompete ACE2 for RBD binding^4,16^. Omicron substitutions may weaken this competition, reducing the ability of such Nbs to prevent receptor engagement unless their binding affinity, kinetic stability, or avidity effects sufficiently compensate.

With the exception of the closely related H11-H4 and H11-D4 pair, the remaining investigated Nbs are sequence-diverse binders rather than variants derived from a common parental Nb; across the panel, CDR sequences and lengths vary most strongly in CDR3. Despite this diversity, our data indicate that, among the SMD-tested Nb interfaces on Omicron BA.2, greater mechanical resistance to unbinding is associated with persistent hydrophobic anchoring at RBM patches near V445–F456 and F490–Y501, little or no mutation-adjacent repulsion, and, in some cases, an acidic CDR residue that forms a stabilizing interaction with Q493R. Within this dataset, NM1230 and WNb-2 highlight complementary design features: NM1230 exhibited the highest mean unbinding work among the Nbs and strong Q493R-centered salt-bridge interactions, whereas WNb-2 showed the highest covariance score, the largest attractive-interaction network, and only one repulsive interaction above the reporting threshold. These results suggest that future CDR engineering should combine aromatic/hydrophobic RBM anchoring with selective electrostatic stabilization near R493 while avoiding polarity or charge mismatches near E484A, Q493R, and Q498R.

Taken together, this work provides a coherent mechanistic picture of Omicron escape across diverse Nb epitopes: Omicron substitutions remodel binding by (i) shifting and broadening bound pose ensembles, (ii) preserving a small set of recurrent hydrophobic RBM anchors, and (iii) selectively introducing repulsive interactions and dynamically anticorrelated motions that erode net interfacial stability. More broadly, our integrated workflow converts long MD trajectories into compact, interpretable “interaction fingerprints” and pathway-resolved descriptions of unbinding thereby pinpointing both what to preserve (shared anchor patches) and what to fix (mutation-induced clash/repulsion sites), offering actionable guidance for structure-guided CDR engineering toward broader variant robustness. By explicitly mapping where Omicron turns attraction into repulsion and how remodeled interfaces fail under force, this study lays a practical foundation for designing next-generation Nbs that remain effective as SARS-CoV-2 continues to evolve.

## Acknowledgement

This work used resources, services and support provided via the COVID-19 HPC Consortium (https://covid19-hpcconsortium.org/) and the Extreme Science and Engineering Discovery Environment (XSEDE), which is supported by National Science Foundation grant number ACI- 1548562. M.Gu. also acknowledges start-up funding by the University of Pittsburgh School of Medicine (UPSOM).

