## Supplementary Material for "From Attraction to Repulsion: Omicron-Driven Rewiring of Nanobody Interfaces on the SARS-CoV-2 RBD"

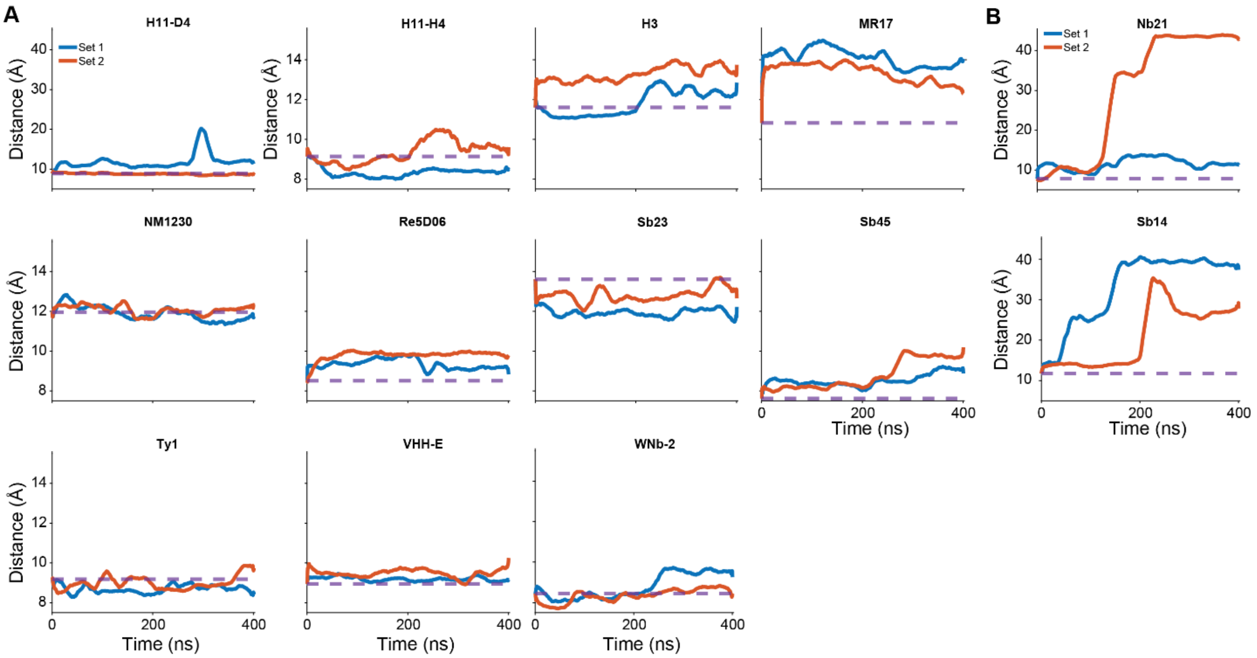


***Figure S1. Nb–RBD distances in Omicron spike MD simulations.*** *RBD residues within 6 Å of the Nb in the Nb–WT spike complexes were identified, and the distance between the center-of-mass (COM) of the Nb CDRs and the COM of these RBD residues was computed across the Nb–Omicron spike trajectories. A horizontal line marks the distance observed in the Nb–WT RBD complexes for reference. (A)*

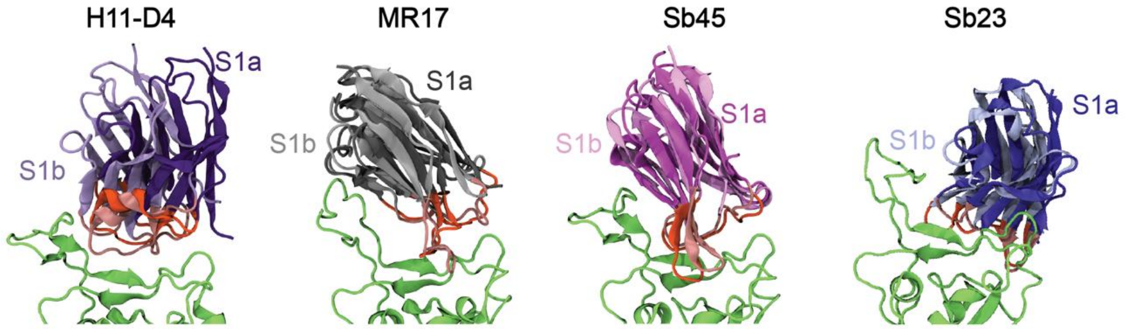


***Figure S2. Superposition of the S1a and S1b binding substates.*** *Representative conformations corresponding to the two closely related low-free-energy substates within basin S1 (S1a and S1b) were extracted from the Nb bound Omicron RBD MD trajectories based on the PCA-defined free-energy landscape. Structures were aligned by least-squares fitting of the RBD $C_{\alpha}$ atoms to the wild-type (WT) RBD reference, and the resulting Nb poses were superimposed to visualize substate-dependent reorientation relative to the RBD. The RBD is shown as a common reference, while the S1a and S1b Nb conformations are displayed in distinct colors as indicated. The comparison highlights a modest change in Nb orientation between substates while maintaining a similar antigen-contact region mediated by the CDRs.*

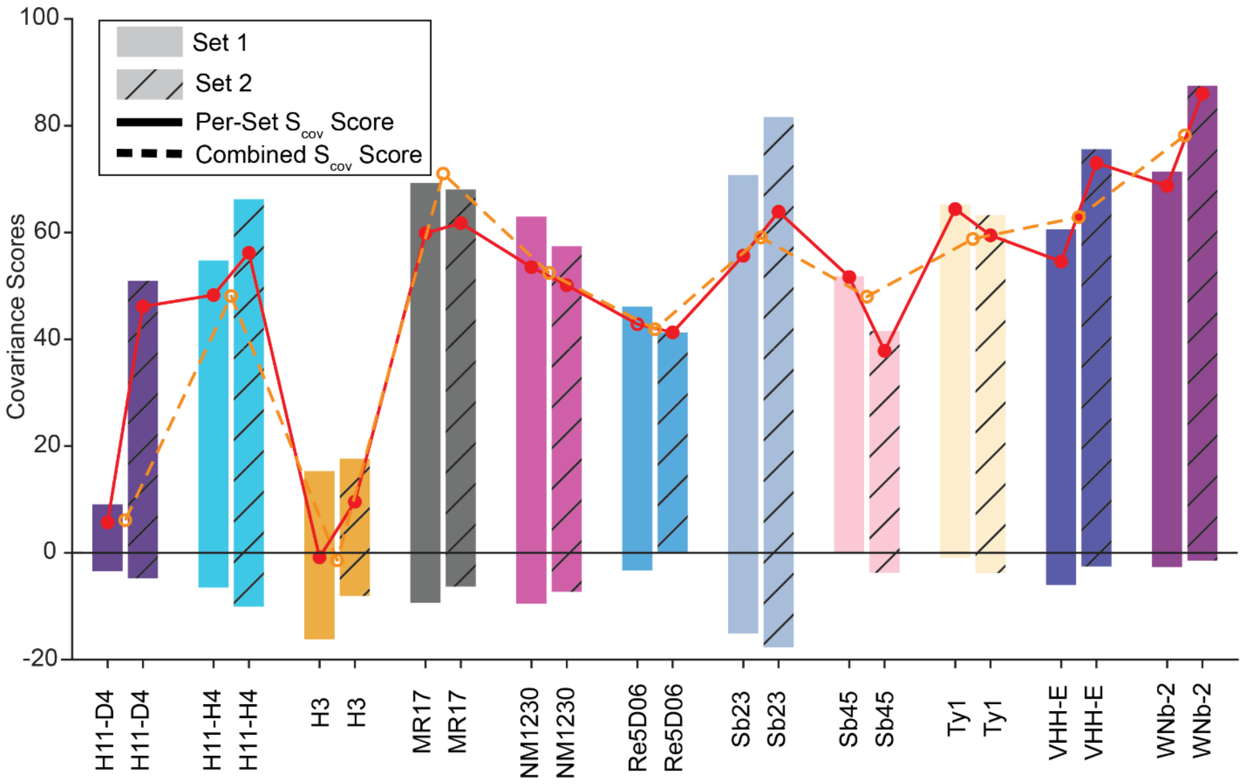


***Figure S3. Cumulative Covariance Scores ($S_{cov}$) Between Nb and RBD residues, per independent trajectory.*** *Each bar represents the positive and negative covariance for the Set 1 (solid bar) and Set 2 (textured bar) replicates. Positive and negative scores are determined by summing up the covariances of each residue's attractive and repulsive scores, respectively. Total scores ($S_{cov}$) were obtained by summing positive and negative covariance scores, for each independent set. These scores indicate robustness in the reported covariance differences between independent trajectories.*

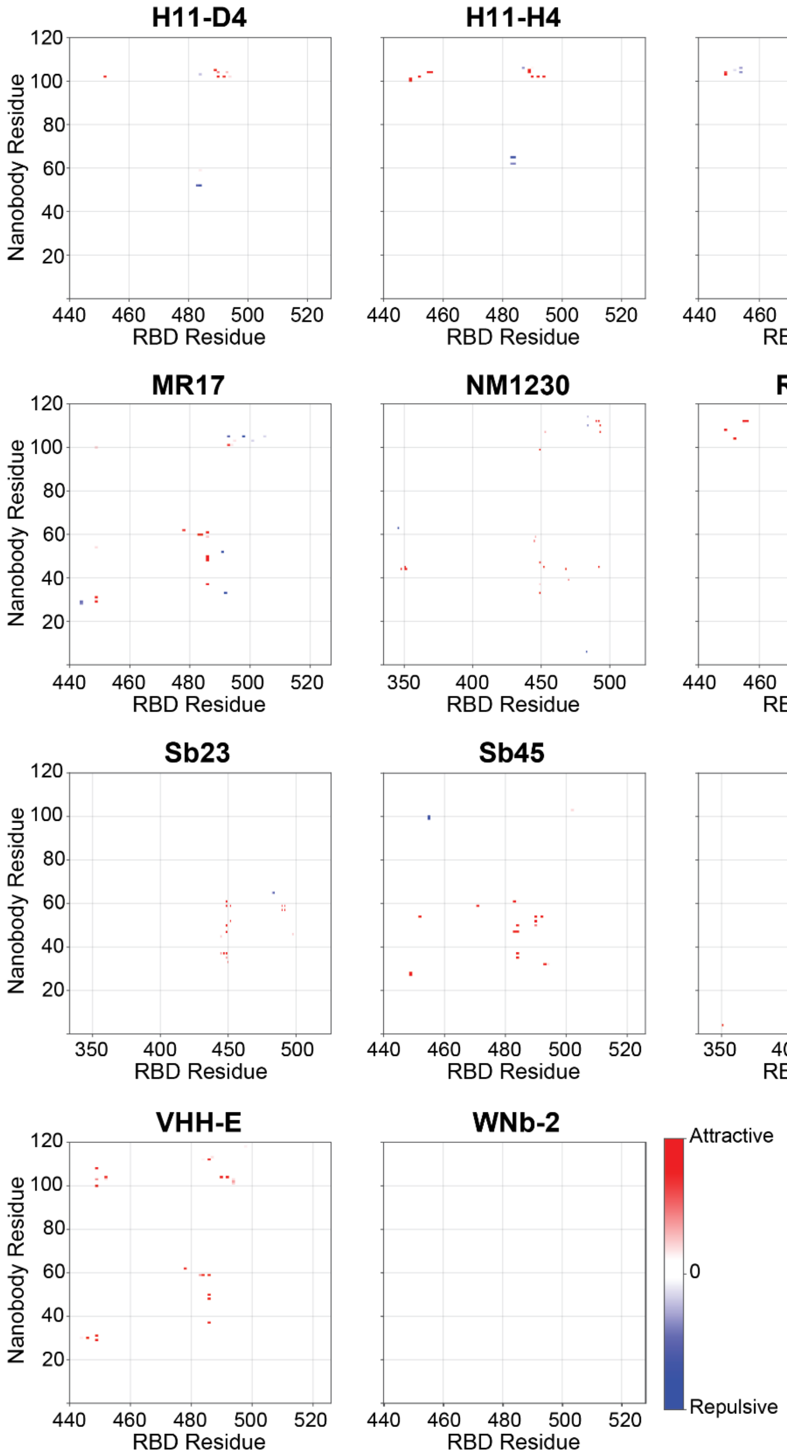


***Figure S4. Interaction frequency and structural mapping for nanobody–Omicron RBD interfaces.*** *Interaction-frequency maps showing the percentage of simulation frames in which each nanobody residue–RBD residue pair satisfies the interaction criteria. The color scale ranges from 0 to +100% (red) for attractive interactions and from 0 to −100% (blue) for repulsive interactions. H11-D4, H11-H4, H3, MR17, NM1230, Re5D06, Sb23, Sb45, Ty1, VHH-E, and WNb-2.*

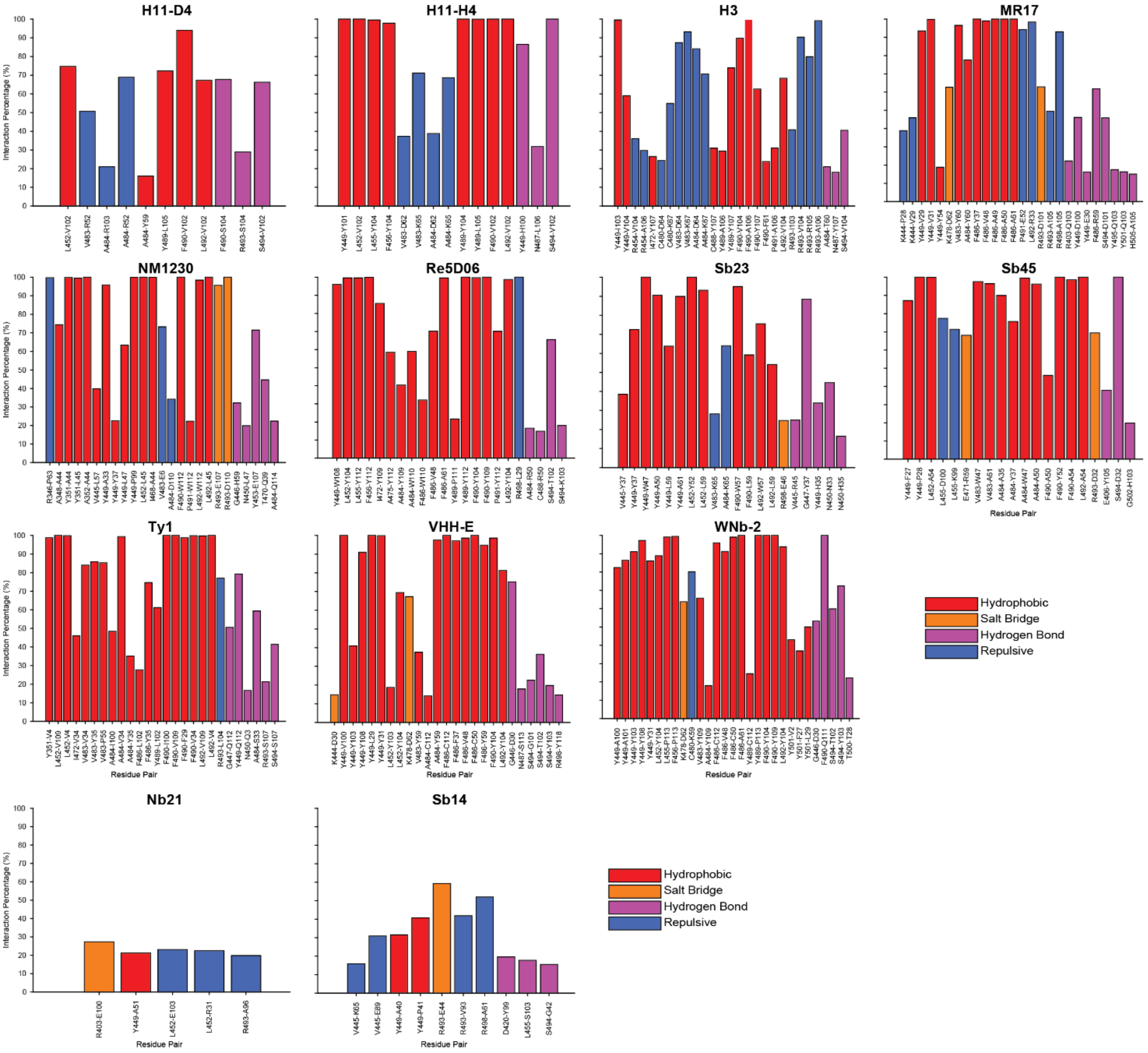


***Figure S5. Interactions Percentages for Nbs.*** *(A) Interaction frequencies of specific residue pairs identified from molecular dynamics simulations for the indicated Nbs (H11-D4, H11-H4, H3, MR17, NM1230, Re5D06, Sb23, Sb45, Ty1, VHH-E, and WNb-2). Interactions are classified and color-coded as hydrophobic (red), salt bridge (orange), hydrogen bond (purple), or repulsive (blue). (B) Corresponding interaction analysis for Nb21 and Sb14.*

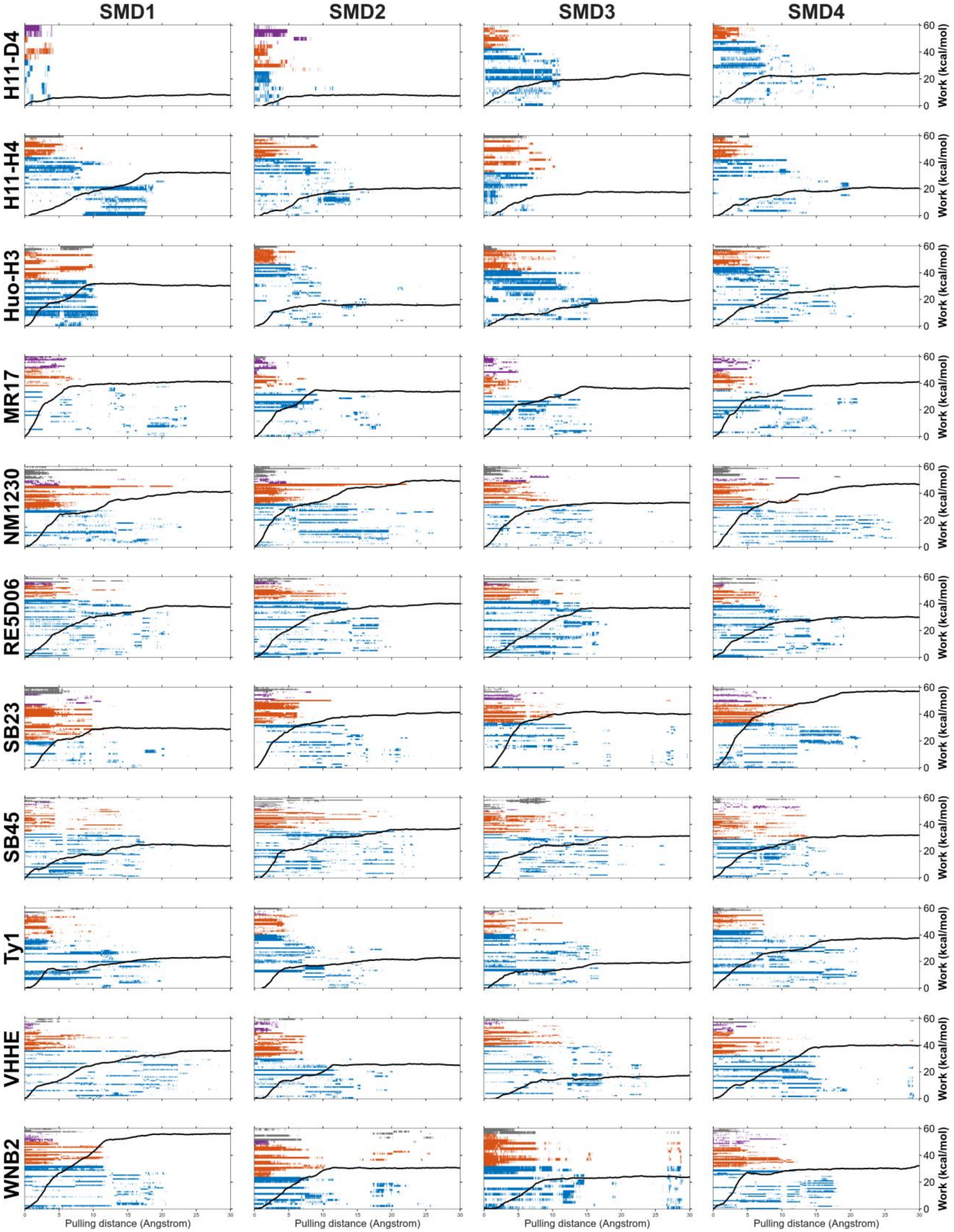


***Figure S6. RBR anchoring correlates with increases in work, during Nb unbinding events.*** *Across all Nbs, re-capture by the RBR is the last stage in the unbinding process, where work spikes*

*and Nbs attempt to recapture the RBD through this flexible ridge.* ***.*** *RBD-Nb interactions are plotted according to the RBD region they bind. Within each RBD region (loop, flat region, RBR, and outer region), pairs are ordered by increasing RBD residue numbering. Interactions between the RBD and Nbs were calculated. Receptor binding ridge (RBR; blue), Flat Region (red), S443-V450 Loop (purple) and outer regions (grey). Overlayed is the work value (kcal mol$^{-1}$) against the pulling direction during SMD simulation, as line plots for each replica SMD simulation, on the right y-axis.*

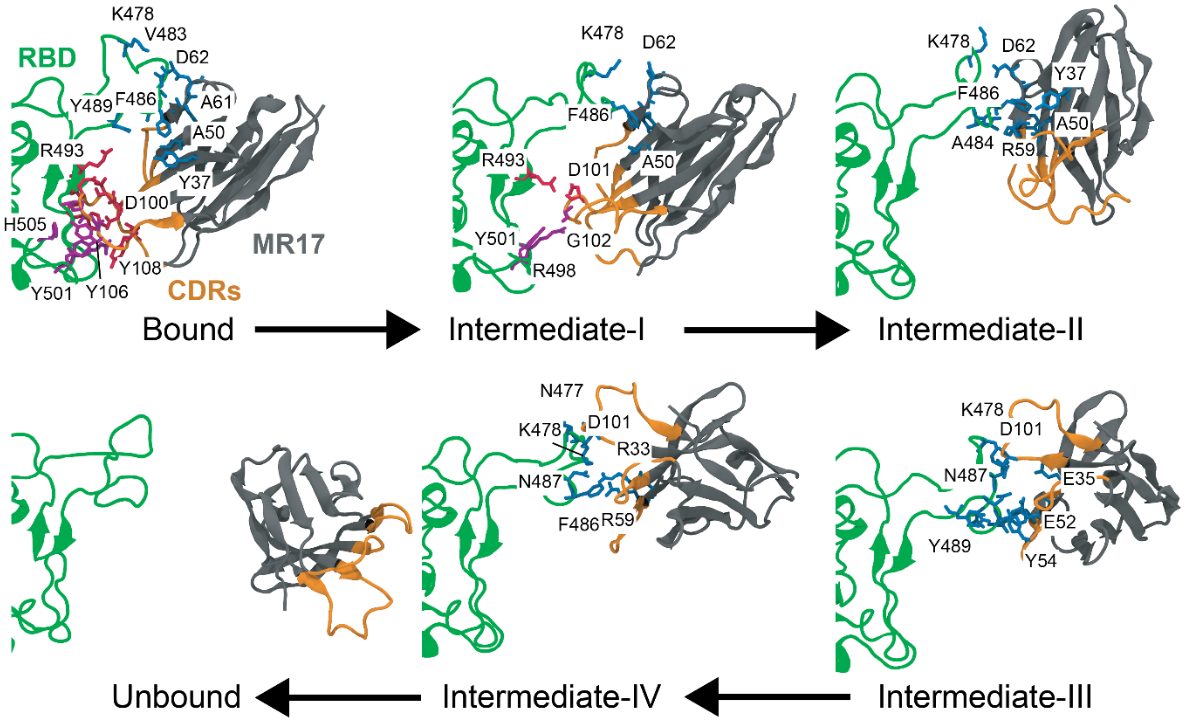


***Figure S7. Residue-level interaction details of the representative MR17 stepwise unbinding pathway.*** *Structural snapshots correspond to the same representative SMD trajectory and conformational states shown in Figure 6A, including the initial bound state, intermediates I–IV, and the fully unbound state. The Omicron RBD is shown in green and MR17 in gray, with the CDR loops highlighted in orange. Interfacial residues participating in the interactions of each conformational state are shown as licorice and labeled.*

***Table S1. Omicron-driven deviation of Nb binding poses from the WT reference*** *Mean ± standard deviation of the nanobody $C_\alpha$ RMSD (Å) relative to the WT Nb–RBD complex, computed over the Omicron MD simulation trajectories after aligning each frame on the RBD. Larger mean RMSD indicates a greater shift from the WT binding pose, whereas larger standard deviation indicates increased pose heterogeneity (broader binding minima / multiple basins). Cell colors summarize the binding-pose classes from the PCA free-energy landscapes (Figure 2): yellow indicates single-basin pose-preserving (high similarity to WT); cyan indicates single-basin with intermediate similarity; magenta indicates single-basin with low similarity; red indicates multi-basin (pose-plastic) sampling with low similarity to WT.*

| Nanobody | Mean | Standard Deviation |
|---|---|---|
| **H11-D4** | 8.2 | 5.5 |
| **H11-H4** | 4.1 | 1.5 |
| **H3** | 7.9 | 4.5 |
| **MR17** | 6.6 | 2.28 |
| **Nb21** | 27.9 | 15.5 |
| **NM1230** | 2.9 | 0.6 |
| **Re5D06** | 6.5 | 1.3 |
| **Sb14** | 18. 6 | 10.7 |
| **Sb23** | 4.0 | 1.1 |
| **Sb45** | 2.8 | 0.9 |
| **Ty1** | 3.1 | 0.6 |
| **VHH-E** | 2.5 | 0.9 |
| **WNb-2** | 3.6 | 1.3 |

***Table S2. Cumulative Covariance Scores ($S_{cov}$) Between Nb and RBD residues.*** *Scores are determined by summing up the covariances of each residue's attractive and repulsive scores. Overall total scores were taken by adding positive and negative covariance scores.*

| Nanobody | Positive | Negative | Total |
|---|---|---|---|
| **H11-D4** | 13.8 | -7.4 | 6.5 |
| **H11-H4** | 68.8 | -22.7 | 46.1 |
| **H3** | 10.4 | -13.5 | -3.1 |
| **MR17** | 76.5 | -5.5 | 71.0 |
| **NM1230** | 58.3 | -8.0 | 50.3 |
| **Re5D06** | 41.3 | -1.7 | 39.5 |
| **Sb23** | 66.8 | -10.1 | 56.7 |
| **Sb45** | 46.2 | -1.4 | 44.8 |
| **Ty1** | 58.3 | -2.6 | 55.7 |
| **VHH-E** | 63.9 | -2.0 | 61.9 |
| **WNb-2** | 76.4 | -0.9 | 75.5 |

*Table S3. Residues Engaging in Multiple Interaction Types Across Simulations*

| Nb | Flagged Residue | RBD Residue | Nb Residue | Interaction Type | Interaction Percentage |
|---|---|---|---|---|---|
| H11-D4 | A484 | A484 | Y59 | HP | 16.1 |
| | | A484 | R103 | R | 21.0 |
| | | A484 | R52 | R | 68.9 |
| | F490 | F490 | S104 | HB | 67.7 |
| | | F490 | V102 | HP | 94.0 |
| | V102 | S494 | V102 | HB | 66.2 |
| | | F490 | V102 | HP | 94.0 |
| | | L452 | V102 | HP | 74.7 |
| | | L492 | V102 | HP | 67.3 |
| H11-H4 | Y449 | Y449 | H100 | HB | 86.5 |
| | | Y449 | Y101 | HP | 100.0 |
| | V102 | S494 | V102 | HB | 100.0 |
| | | F490 | V102 | HP | 100.0 |
| | | L452 | V102 | HP | 100.0 |
| | | L492 | V102 | HP | 100.0 |
| H3 | A484 | A484 | T60 | HB | 21.0 |
| | | A484 | D64 | R | 84.0 |
| | | A484 | K67 | R | 70.6 |
| | A106 | F490 | A106 | HP | 99.6 |
| | | P491 | A106 | HP | 31.0 |
| | | Y489 | A106 | HP | 29.4 |
| | | R454 | A106 | R | 29.7 |
| | | R493 | A106 | R | 99.1 |
| | I103 | Y449 | I103 | HP | 99.6 |
| | | R493 | I103 | R | 40.8 |
| | V104 | S494 | V104 | HB | 40.5 |
| | | F490 | V104 | HP | 89.8 |
| | | L492 | V104 | HP | 68.3 |
| | | Y449 | V104 | HP | 59.0 |
| | | R454 | V104 | R | 36.0 |
| | | R493 | V104 | R | 90.3 |
| | Y107 | N487 | Y107 | HB | 18.1 |
| | | C488 | Y107 | HP | 30.9 |
| | | F490 | Y107 | HP | 62.6 |
| | | I472 | Y107 | HP | 26.5 |

| | | | | | |
|---|---|---|---|---|---|
| | | Y489 | Y107 | HP | 73.8 |
| MR17 | R493 | R493 | D101 | SB | 62.8 |
| | | R493 | A105 | R | 49.4 |
| | A105 | H505 | A105 | HB | 15.1 |
| | | R493 | A105 | R | 49.4 |
| | | R498 | A105 | R | 93.0 |
| | V29 | Y449 | V29 | HP | 93.4 |
| | | K444 | V29 | R | 45.9 |
| NM1230 | A484 | A484 | Q114 | HB | 22.4 |
| | | A484 | D110 | R | 34.2 |
| | D110 | R493 | D110 | SB | 100.0 |
| | | A484 | D110 | R | 34.2 |
| | L47 | N450 | L47 | HB | 19.9 |
| | | Y449 | L47 | HP | 63.3 |
| Sb23 | Y449 | Y449 | H35 | HB | 33.9 |
| | | Y449 | A50 | HP | 90.5 |
| | | Y449 | A61 | HP | 89.9 |
| | | Y449 | L59 | HP | 63.8 |
| | | Y449 | W47 | HP | 100.0 |
| | | Y449 | Y37 | HP | 72.5 |
| | Y37 | G447 | Y37 | HB | 88.4 |
| | | V445 | Y37 | HP | 38.5 |
| | | Y449 | Y37 | HP | 72.5 |
| Ty1 | A484 | A484 | S33 | HB | 59.3 |
| | | A484 | I100 | HP | 48.4 |
| | | A484 | V34 | HP | 99.4 |
| | | A484 | Y35 | HP | 35.2 |
| VHH-E | Y103 | S494 | Y103 | HB | 19.7 |
| | | L452 | Y103 | HP | 18.7 |
| | | Y449 | Y103 | HP | 40.9 |
| WNb-2 | F490 | F490 | Q111 | HB | 100.0 |
| | | F490 | Y104 | HP | 100.0 |
| | | F490 | Y109 | HP | 100.0 |
| | Y103 | S494 | Y103 | HB | 72.5 |
| | | Y449 | Y103 | HP | 91.1 |

**Simulation Information**

| Nanobody | cMD (ns) | SMD (ns) |
|---|---|---|
| **H11-D4** | (a) 400, (b) 400 | (a1) 300, (a2) 300, (b1) 300, (b2) 300 |
| **H11-H4** | (a) 400, (b) 400 | (a1) 300, (a2) 300, (b1) 300, (b2) 300 |
| **H3** | (a) 400, (b) 400 | (a1) 300, (a2) 300, (b1) 300, (b2) 300 |
| **MR17** | (a) 400, (b) 400 | (a1) 300, (a2) 300, (b1) 300, (b2) 300 |
| **Nb21** | (a) 400, (b) 400 | - |
| **NM1230** | (a) 400, (b) 400 | (a1) 300, (a2) 300, (b1) 300, (b2) 300 |
| **Re5D06** | (a) 400, (b) 400 | (a1) 300, (a2) 300, (b1) 300, (b2) 300 |
| **Sb14** | (a) 400, (b) 400 | - |
| **Sb23** | (a) 400, (b) 400 | (a1) 300, (a2) 300, (b1) 300, (b2) 300 |
| **Sb45** | (a) 400, (b) 400 | (a1) 300, (a2) 300, (b1) 300, (b2) 300 |
| **Ty1** | (a) 400, (b) 400 | (a1) 300, (a2) 300, (b1) 300, (b2) 300 |
| **VHH-E** | (a) 400, (b) 400 | (a1) 300, (a2) 300, (b1) 300, (b2) 300 |
| **WNb-2** | (a) 400, (b) 400 | (a1) 300, (a2) 300, (b1) 300, (b2) 300 |

**Detailed Interaction Information on Nb-Omicron RBD Interactions**

Hydrophobic interactions, hydrogen bonds, salt bridges, and repulsive interactions were abbreviated as HP, HB, SB, and R, respectively. High-frequency (>49%) and medium-to-high–frequency (>14%) interactions for the analyzed nanobodies (Nbs) were reported for both attractive and repulsive interactions, respectively.

**H11-D4:** H11-D4 exhibits weak yet spatially localized correlated motions with the RBD, characterized by sparse positive covariance centered on residues Y449–Y453 and C488–Y495, along with limited anticorrelated features near the V483–A484 region, yielding an overall interaction score of 6.5. These dynamics are indicative of a shallow interaction footprint, comprising 4 HP, 2 HB and 3 R. The HP localize primarily to residues L452, Y489–F490, and L492, while HB involve residues F490 and S494; the repulsive interactions are confined to the V483–A484 region.

**H11-H4:** H11-H4 showed predominantly positive correlation with RBD segments Y449–Y453, G447–R457, S469–Y473, and C488–G496, while exhibiting anticorrelated behavior across P479–N487 giving an interaction score of 46.1. These dynamics map to an interface with attractive contacts (HP/HB) concentrated at residues Y449–F456 and Y489–S494, countered by repulsive (aliphatic–polar) interactions localized to residues V483–A484. Overall, the interface is HP-dominated (8 HP, 2 HB, 4 R), and notably, half of the repulsive contacts arise from persistent aliphatic–charged clashes/repulsion between the Omicron E484A site and H11-H4 residues D62 and K65 (39% and 69% occupancy, respectively), highlighting how this binder accommodates Omicron by forming new unfavorable contacts, distinct from the charge–charge repulsion previously observed[31] for the Beta variant.

**H3:** H3 exhibited a mixed covariance signature with positive correlations at G447–Y449, I468–Y473, and Y489–G496 opposed by negative correlations at L452–Y453 and C480–N487, leading to a slightly negative interaction score (–3.1) that matches its interaction balance: attractive contacts are primarily hydrophobic (Y449, Y489–F490, and L492), but they are likely outweighed by extensive repulsive interactions (aliphatic–polar and charge–charge) at R454, C480–A484, and R493. Consistent with this, H3 has the highest repulsion burden (7 HP, 1 HB, 12 R), including six repulsive interactions involving Omicron-specific mutation sites (Q493R and E484A), all with occupancies between 40-99%.

**MR17:** MR17 showed correlated motion with V445–Y453 and I472–R498 but anticorrelated coupling across F490–H505 (interaction score 71.0), consistent with a contact pattern in which attractive interactions (HP/SB/HB) are concentrated at residue Y449 and across K478–R493, while repulsive (aliphatic–polar) interactions appear at residues K444, P491, L492, R493, and R498. The interaction inventory comprises 9 HP, 2 SB, and 6 R interactions, with most repulsive contacts occurring within the Nb–RBD core and half exhibiting >93% occupancy; additionally, MR17 forms two SB interactions involving the T478K and Q493R mutation in the RBM. Notably, R493 forms a SB with D101 at 62% occupancy, while also showing a repulsive interaction with A105 at 49% frequency.

**NM1230:** NM1230 presented positive correlations spanning A348–N354, G446–L452, E465–E471, and C488–F497 with negative correlations at G482–N487, giving an interaction score of 50.3; this aligns with an attractive interaction footprint at A348–A352, V445–L452, I468, and F490–L492, accompanied by repulsive interactions (aliphatic-polar) at residues R346, V483 and A484. The overall interaction composition is 12 HP, 2 SB, 2 HB, and 3 R, with one repulsive interaction occurring within the RBM/Nb–RBD core; importantly, NM1230's extended attractive (HP) contacts are highly persistent (>99% occupancy) while there is a singular extended repulsive contact, (~99% occupancy). Additionally, NM1230 forms strong SB interactions with the Q493R mutation (95.6% and 100% occupancy) plus a shorter repulsive interaction with E484A (34.2%), which is also noted as its lowest repulsive occupancy interaction.

**Re5D06:** Re5D06 exhibited positive correlations along G446–R457 and A484–F497 with localized negative correlations at I468–E471 and G482–Y489 (interaction score 39.5), and it engages the RBM through an attractive footprint spanning residues Y449–F456, I472–A475, and A484–L492; in the interaction tally this manifests as a largely hydrophobic interface (14 HP) supported by a single HB (1 HB) and a single repulsive interaction (1 R). This single repulsive interaction is fully persistent and occurs between the omicron mutation site Q498R. Re5D06 also maintains an HP involving the Omicron-specific E484A site (59% occupancy) and its W110.

**Sb23:** Sb23 showed positive correlations at A344–V350, K440–Y453, and P491–T500 with negative correlations at I468–E471 and G482–Y489 (interaction score 56.7), and it forms attractive contacts (HP/SB/HB) primarily across residues V445–L452 and F490–L492, counterbalanced by repulsive (aliphatic–polar) interactions at residues V483–A484. In the interaction tally, Sb23 comprises 11 HP, 1 HB, and 2 R interactions that are described as being confined to the Nb–RBD core, and it engages with Omicron mutation sites via an SB interaction a repulsive interaction with E484A (64.0%).

**Sb45:** Sb45 showed positive correlations across G447–L455, I468–I472, and N481–F497 (score 44.8), and it forms an attractive network (HP/SB/HB) at residues Y449, L452, E471, V483–A484, and F490–R493, offset by two repulsive (aliphatic–polar) interactions at residue L455. In total, Sb45 forms 12 HP, 2 SB, and 2 R interactions, and 5 of its 14 attractive interactions occur at Omicron mutation sites (4 HP interactions with A484 and 1 SB interaction with R493).

**Ty1:** Ty1 displayed positive correlations across a broad footprint (S349–W353, K444–Y453, I468–I472, and N481–G496) and a minimal repulsive band between N481–V483, giving an interaction score of 55.7, and it forms mainly attractive interactions at residues Y351, G447–L452, I472, and V483–L492. The interaction inventory is almost entirely stabilizing (15 HP and 2 HB), including multiple contacts involving the Omicron E484A site (2 HP and 1 HB). A single repulsive interaction (77.0%) at the Q493R omicron mutation disrupts this mostly hydrophobic code. There are no salt-bridge interactions reported for this binder.

**VHH-E:** VHH-E showed strong positive correlations across S443–Y449, A475–K478, and A484–V503 (interaction score 61.87) and forms an exclusively attractive interaction network (HP/SB/HB) spanning residues K444–L452, K478, and A484–L492; accordingly, its interaction tally includes 13 HP, and 1 SB interactions with no repulsive contacts. Two especially persistent interactions are an HP contact with Omicron mutation sites A484 (97.6% occupancy) and an SB involving K478 (67.3% occupancy).

**WNb-2:** WNb-2 displayed strong correlated coupling across S443–R457 and G482–G502 with a narrow anticorrelated hotspot at G482, yielding the highest interaction score in this set of Nbs (75.5), and it forms extensive attractive contacts (HP/HB/SB) across G446–F456, V483–S494, and R498–Y501; the only unfavorable feature is a single aliphatic–polar repulsive interaction at residue 480 that is persistent (80.14% occupancy). In total, WNb-2 forms 18 HP, 3 HB, 1 SB, and 1 R interactions, including SB involving the Omicron mutation site at Q478R (63.8%), as well as HP contact with N501Y (50.0%).

**Supplementary Videos S1–S12.** Steered molecular dynamics (SMD) simulations of RBD–binder complexes. Each video shows four independent SMD pulling trajectories, with the pulling direction indicated by a red arrow. Work–distance profiles are displayed in the background behind the simulation trajectories. In all videos, RBD is shown in green.

**Supplementary Video S1.** RBD–ACE2 complex; ACE2 is shown in beige.

**Supplementary Video S2.** RBD–H11-D4 complex; H11-D4 is shown in indigo.

**Supplementary Video S3.** RBD–H11-H4 complex; H11-H4 is shown in cyan.

**Supplementary Video S4.** RBD–H3 complex; H3 is shown in orange.

**Supplementary Video S5.** RBD–MR17 complex; MR17 is shown in gray.

**Supplementary Video S6.** RBD–NM1230 complex; NM1230 is shown in magenta.

**Supplementary Video S7.** RBD–Re5D06 complex; Re5D06 is shown in blue.

**Supplementary Video S8.** RBD–Sb23 complex; Sb23 is shown in lilac.

**Supplementary Video S9.** RBD–Sb45 complex; Sb45 is shown in pink.

**Supplementary Video S10.** RBD–Ty1 complex; Ty1 is shown in white.

**Supplementary Video S11.** RBD–VHH-E complex; VHH-E is shown in deep blue.

**Supplementary Video S12.** RBD–WNb-2 complex; WNb-2 is shown in purple.
